# Maturase K forms a plastidial splicing complex with a neofunctionalized branching enzyme

**DOI:** 10.1101/2025.01.08.628318

**Authors:** Yuanyuan Liang, Yang Gao, Andrea Fontana, Melanie R. Abt, Adam P. Gicgier, Chun Liu, Mayank Sharma, Reimo Zoschke, Samuel C. Zeeman, Barbara Pfister

**Affiliations:** Institute of Molecular Plant Biology, ETH Zurich, Universitätstrasse 2, 8092 Zurich, Switzerland; Max Planck Institute of Molecular Plant Physiology, Am Mühlenberg 1, 14476 Potsdam-Golm, Germany; Department of Crop Genetics, John Innes Centre, Norwich, NR4 7UH, UK; Crop Science Centre, University of Cambridge, Lawrence Weaver Rd, Cambridge, CB3 0LE, UK

## Abstract

Chloroplast group IIA introns derive from bacterial ribozymes. Their splicing likely requires Maturase K (MatK), which has been largely inaccessible to functional analyses being itself a chloroplast intron-encoded protein. Here we show that MatK physically interacts with a conserved, essential plastid-localized homolog of starch-branching enzymes (BEs), dubbed MATURASE K INTERACTING PROTEIN1 (MKIP1). We demonstrate that MKIP1 proteins have lost BE activity and acquired an insertion enabling direct interaction with the N-terminal region of MatK. Arabidopsis MKIP1 specifically co-precipitates all known intron targets of MatK. Induced *MKIP1* silencing results in pale newly emerging leaves, in which the splicing of these intron targets is strongly reduced. Our data suggest that MKIP1 functionally diverged from canonical BEs to facilitate splicing in conjunction with MatK. We propose that the N-terminus of MatK, in turn, has evolved from an RNA-binding domain into a platform for protein interaction, helping its transition towards a general splicing factor.

## Introduction

Chloroplast gene expression involves the splicing of ∼20 introns, a process essential for plant viability^1,2^. Most of these belong to group II introns derived from mobile bacterial elements, which are also probable predecessors of nuclear mRNA introns^3^. In bacteria, group II introns were shown to have six domains (denoted as domain I to VI) that fold into a complex tertiary structure^4^. Although splicing is catalyzed by the ribozyme intron RNA itself, efficient splicing usually requires a maturase protein^5^. These are typically encoded within intron domain IV and have an N-terminal reverse transcriptase (RT) domain, followed by a maturase (X) domain, among others^6^. During splicing, the RT domain of the bacterial maturase stably docks the protein onto the intron in a sequence-specific manner, allowing the X domain to make flexible contacts with the intron core^6–8^. Domain X then facilitates critical RNA conformations during splicing by stabilizing the intron’s catalytic core and the position of the bulged adenosine in domain VI - the nucleophile of the first splicing step^9^.

In land plants chloroplasts, only the *trnK* intron encodes a maturase-like protein, Maturase K (MatK). Tobacco MatK was observed to bind to six^10^ or seven^11^ introns, i.e. to all group IIA introns except the divergent^12,13^ introns in *clpP* and (possibly) *trnV*. MatK likely helps the splicing of its binding targets, since splicing is reduced upon MatK reduction by genetic or chemically-induced disruption of plastid translation, and since *matK* gene loss coincides with intron loss in several parasitic plants^14^. These observations imply that MatK has evolved into a general splicing factor, but direct functional analyses of MatK have been hindered by the lack of (hypomorphic) mutants of this likely essential gene^11^.

MatK has retained a conserved X domain, but its RT domain has largely diverged and the N-terminal RT motifs of bacterial maturases are no longer recognizable^15^. MatK also lacks a positively charged N-terminal extension that, together with the N-terminal RT region, enables bacterial maturases to form strong and sequence-specific interactions with intron domain IV^7,8^. The potential loss of specific RNA binding may have helped MatK to act on multiple introns, but likely comes at the cost of a lower RNA affinity and may negatively affect its orientation on the intron^6^. In addition, the plastidial introns have lost RNA-RNA interaction motifs involved in the folding of bacterial introns^16^. These deviations may explain why plastidial group IIA intron splicing requires an array of nuclear-encoded splicing factors, such as CRS1 (Chloroplast RNA Splicing 1), RNC1 and WTF1 (“what’s this factor?”)^17–19^. These possibly assist in the initial folding and stabilization of the precursor^1,2^, but have not been reported to associate directly with MatK.

Here we report a protein that physically interacts with MatK to help plastidial intron splicing in Arabidopsis. This protein was previously named BRANCHING ENZYME1 (*At*BE1) (AT3G20440), based on its homology to starch-branching enzymes (BEs), which introduce α-1,6-linkages into the polymers comprising starch, a plastidial storage carbohydrate^20^. However, in contrast to the other two Arabidopsis BEs (*At*BE2 and *At*BE3), *At*BE1 has no apparent BE activity^21,22^. In addition, Arabidopsis *be1* mutants are embryo defective and inviable^23,24^, while *be2* and *be3* single mutants are phenotypically normal^21^. The *be2/be3* double mutant lacks starch and grows poorly but is viable^21^, as are, to our knowledge, all other Arabidopsis mutants with primary defects in starch metabolism described to date. We show that *At*BE1 is a conserved, neofunctionalized BE that has acquired a new role in plastid intron splicing by forming a splicing complex with *At*MatK. To recognize its function and avoid confusion with canonical BEs, we renamed it to <u>M</u>ATURASE <u>K I</u>NTERACTING <u>P</u>ROTEIN1, or MKIP1.

## Results

### *At*MKIP1 is required for plant viability

We reconfirmed that *At*MKIP1 is essential for embryo development^23,24^ using two lines (*mkip1-1* and *mkip1-2*) carrying T-DNA insertions in introns 15 and 16 of the *AtMKIP1* (AT3G20440) gene, respectively (**Fig. 1a, Supplementary Fig. 1**). No *mkip1^-/-^* mutant plants could be identified from the progeny of mother plants heterozygous for either allele (192 and 100 plants analyzed for *mkip1-1* and *mkip1-2,* respectively). In addition, siliques from heterozygous plants contained ∼25% white seeds with embryos arrested at the heart stage (**Fig. 1a, Supplementary Fig. 1**). These phenotypes could be complemented by *P_UBQ10_*-driven, constitutive expression of the coding sequence (CDS) of *At*MKIP1 fused to a C-terminal mCitrine or eYFP (named YFP hereafter) tag in heterozygous *mkip1-1* or *mkip1-2* plants. Complemented *mkip1-1 ^-/-^* and *mkip1-2^-/-^* plants appeared wild-type (WT) like although they grew slightly slower (**Fig. 1, Supplementary** Figs. 1, 2c). Confocal microscopy of *P_35S_*-driven, stable expression of *At*MKIP1-YFP in WT Arabidopsis confirmed that *At*MKIP1 localizes to the chloroplasts^23^ of epidermal cells in a fairly homogeneous manner (**Fig. 1b**).

**Fig. 1.**
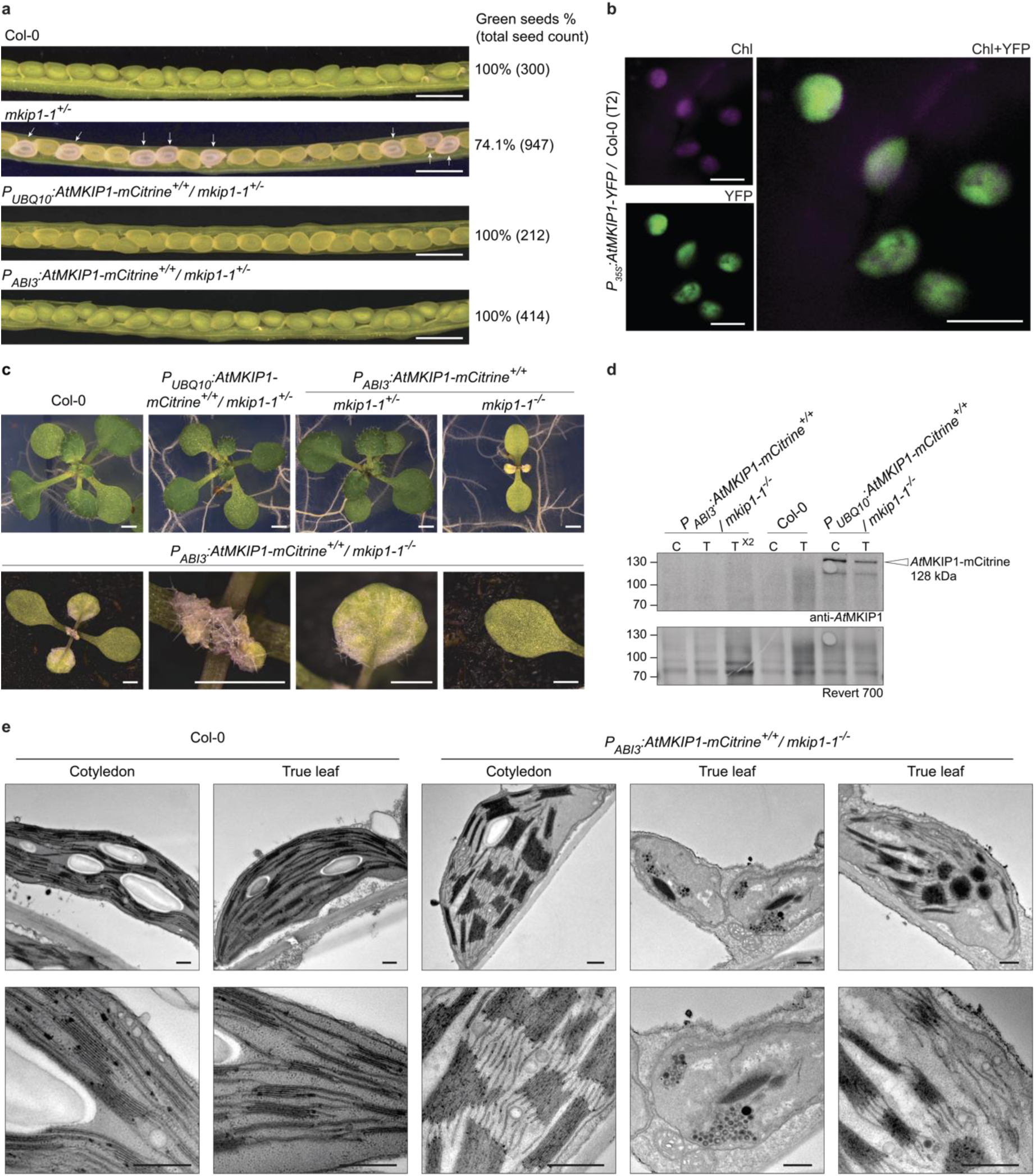
*At*MKIP1 is essential for embryonic and post-embryonic development. **a** Opened siliques of wild type (Col-0), *mkip-1^+/-^*, and *mkip-1^+/-^* plants transformed with *P_UBQ10_:AtMKIP1-mCitrine* or *P_ABI3_:AtMKIP1-mCitrine* constructs. White seeds are indicated by arrows. Percentages indicate the proportion of green seeds among the total seeds (number of seeds analyzed in parentheses). Scale bars: 1 mm. **b** Confocal microscopy of chlorophyll autofluorescence (Chl, magenta) and YFP signal (YFP, green) of *At*MKIP1-YFP in Arabidopsis leaf epidermis. Two other independent lines gave similar results. Scale bars: 5 µm. **c** *mkip-1^-/-^* seedlings rescued with the *P_ABI3_:AtMKIP1-mCitrine* construct. Top panel: Photographs of representative 16-day-old seedlings of the indicated genotypes grown on 1/2-strength MS agar plates. Bottom panel: Micrographs of a representative 4-week-old, soil-grown *P_ABI3_:AtMKIP1-mCitrine^+/+^/ mkip-1^-/-^* plant showing the whole plant, the meristem, a true leaf, and a cotyledon (from left to right). Scale bars: 1 mm. Equivalent analyses in the *mkip-2^-/-^*background yielded similar results (**Supplementary Fig. 1**). **d** Anti-*At*MKIP1 immunoblot of total proteins from cotyledon (C) or true leaf (T) tissue of the indicated lines. Samples were loaded on an equal fresh weight basis. X2 indicates double loading. The predicted molecular weight of *At*MKIP1-mCitrine refers to the mature proteoform without the chloroplast transit peptide. Native *At*MKIP1 (98 kDa) in Col-0 could not be detected due to insufficient antibody sensitivity. Revert 700 is a total protein stain (loading control). **e** Transmission electron micrographs of chloroplasts from 4-week-old, soil-grown Col-0 and *mkip-1^-/-^* plants rescued with the *P_ABI3_:AtMKIP1-mCitrine* construct. Two additional replicate plants and equivalent analyses in the *mkip-2^-/-^* background gave similar results. Scale bars: 0.5 µm.

In an attempt to bypass the embryonic arrest of *mkip1^-/-^* mutants, we transformed heterozygous *mkip1-1^+/-^* and *mkip1-2^+/-^* plants with a construct expressing *At*MKIP1-mCitrine under the control of the embryo-specific *ABI3* promoter^25^. Siliques from *mkip1^+/-^* plants homozygous for the construct did not contain any white seeds, indicating a successful rescue of the embryo-defectiveness of *mkip1^-/-^* (**Fig. 1a, Supplementary Fig. 1d**). We recovered rescued seedlings in both *mkip1-1^-/-^* and *mkip1-2^-/-^*backgrounds at the expected frequency of ∼25%. However, after germination, these seedlings developed serrated, white true leaves and abnormal meristems and were seedling lethal (**Fig. 1c, Supplementary Fig. 1e**). These phenotypes were similar to *P_ABI3_*-rescue lines of mutants severely impaired in chloroplast differentiation due to primary defects in plastidial translation or splicing^26^. Seedling lethality of rescued *mkip1-1^-/-^* could not be rescued by growing plants on ½-strength MS medium supplemented with 1% sucrose (**Supplementary Fig. 1c**).

Plastids from true leaves of *P_ABI3_*-rescued *mkip1-1^-/-^*seedlings mostly contained collapsed thylakoid stacks and accumulated electron-dense, vesicular structures of unknown nature. Occasionally, we observed chloroplast-like plastids with large condensed thylakoid stacks (**Fig. 1e**, outer right micrograph), reminiscent of structures observed upon co-suppression of the plastidial ribosomal protein *At*bS1c (RPS1)^27^. The cotyledons were pale green, and their plastids had abnormally stacked thylakoids (**Fig. 1c, e**). Presumably, the cotyledons were less affected due to residual *At*MKIP1-mCitrine protein remaining in cotyledon tissue from embryogenesis, although *At*MKIP1-mCitrine levels were below the detection limit (**Fig. 1d**). These phenotypes strongly suggest that *At*MKIP1 is essential for plant viability also after embryogenesis and are consistent with a putative essential role of *At*MKIP1 in chloroplast function, but not in starch metabolism.

### *At*MKIP1 and orthologs have lost branching enzyme activity

Phylogenetic analyses showed that the canonical BEs involved in starch biosynthesis of land plants and streptophyte algae belong to either class I (which is absent in Arabidopsis) or class II (to which *At*BE2 and *At*BE3 belong; **Fig. 2a**), confirming previous observations^28^. The BEs of chlorophyte algae also mostly belong to these classes, whereas those of glaucophyte and rhodophyte algae, which produce floridean starch in the cytosol^29^, form a separate group. *At*MKIP1 and its orthologs form yet another, well-separated clade of class III BEs^28^, which share the structural features of *At*MKIP1 described below. We identified at least one class III BE in each photoautotrophic land plant examined and in some streptophyte algae, but not in any other alga (**Fig. 2a, Supplementary Fig. 3a**), suggesting that it arose during the colonization of land by plants (**Fig. 2a**, **Supplementary Fig. 3a**). The origin of class III BEs is unclear, but their distinct exon-intron gene organization^28^ (**Supplementary Fig. 3b**), renders it unlikely that they arose from the duplication of a class I or II BE gene.

**Fig. 2.**
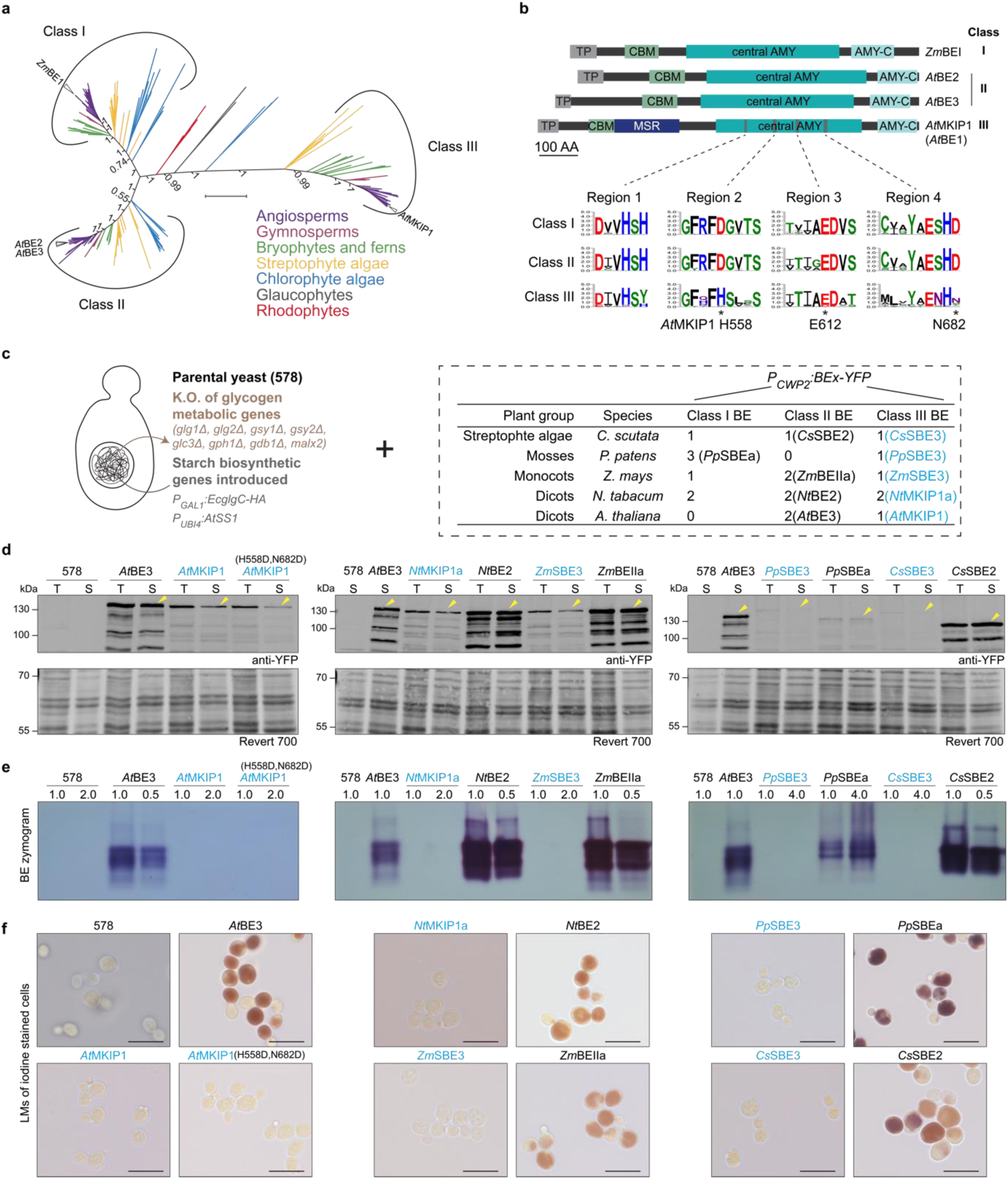
*At*MKIP1 and orthologous proteins are non-canonical, catalytically inactive branching enzymes (BEs). **a** Unrooted Bayesian phylogenetic tree of 254 BE proteins from 85 algal and land plant species (listed in **Supplementary Fig. 3a**) colored according to their phylogenetic groups. Relevant posterior branch probabilities are indicated next to the branches. Scale bar: 1 amino acid substitution per site. **b** Top panel: Schematic representation of Arabidopsis BEs from different classes. Maize BEI (*Zm*BEI) is included as a representative of class I BE, which is absent in Arabidopsis. Boxes depict the chloroplast transit peptide (cTP, *gray*), the carbohydrate-binding module (CBM, *green*), the MKIP1-specific region (MSR, *dark blue*), the central α-amylase domain (central AMY, *cyan*), and the C-terminal α-amylase domain (AMY-C, *light cyan*). Lower panel: WebLogos of protein motifs highly conserved in class I and II BEs, based on the sequence alignment used for the phylogenetic tree in (**a**). Asterisks mark the catalytic triad. **c** Genetic modifications of yeast strain “578”, enabling it to produce starch-like glucans upon introduction of an active BE. The box lists the plant BEs expressed in “578” yeast together with the numbers of BEs present in the respective species. Class III BEs are highlighted in blue. BEs were stably expressed from the yeast nuclear genome via the strong constitutive *CPW2* promoter and fused to a C-terminal YFP tag. **d** Immunoblot detection of YFP-tagged BEs in total (T) or soluble (S) yeast protein extracts. Yellow arrows mark the protein bands matching the proteins’ molecular weights. Revert 700, total protein stain. **e** BE activity gels (zymograms) of yeast soluble protein extracts. Native gels containing glycogen were incubated in the presence of glycogen phosphorylase, the activity of which is stimulated by the presence of an active BE. Glucan products were visualized by iodine staining of the gel. Numbers indicate the relative protein load. **f** Light micrographs of yeast cells stained with iodine. Dark brown/purple staining indicates the presence of glucans. Scale bars: 10 µm.

Class III BEs retain a classical BE domain architecture consisting of a family 48 carbohydrate-binding module (CBM) and α-amylase domains (**Fig. 2b**). Nonetheless, they display striking differences from canonical BEs, with which they share only ∼30% amino acid (AA) identity^28^. First, class III BEs possess an MKIP1-specific region (MSR) of ∼150 AA after the CBM. This stretch is enriched for positively and negatively charged AA (especially lysines and glutamates, respectively) and is predicted to form a largely disordered region protruding from the *At*MKIP1 core (**Supplementary Fig. 4**). An *At*MKIP1(ΔMSR)-YFP version lacking the MSR failed to complement the embryo defects of *mkip1-1^-/-^*, despite being expressed as a soluble protein, indicating a key role for *At*MKIP1 function (**Supplementary Fig. 5**).

Class III BEs also carry substitutions in several residues required for starch branching catalysis, including two aspartate residues of the catalytic triad^30^ (H558 and N682 in *At*MKIP1; **Fig. 2b**), consistent with *At*MKIP1’s apparent absence of BE activity^21,22^. To test whether glucan BE activity is also absent in orthologous proteins, we expressed MKIP1 versions from tobacco (*Nicotiana tabacum*), maize (*Zea mays*), a moss (*Physcomitrium patens*) and a streptophyte alga (*Coleochaete scutata*) as YFP-tagged proteins in the yeast *Saccharomyces cerevisiae* (**Fig. 2c**). The parental yeast strain used, “578”, contains all components required for glucan production except a BE, as it has non-yeast genes encoding a starch synthase (*At*SS1) and an enzyme producing its substrate ADPglucose (*Ec*glgC-TM-HA), but lacks its endogenous glycogen BE as well as all other yeast glycogen-metabolic genes^31^. All class III BEs were expressed as soluble proteins in yeast, although in some cases at low levels (**Fig. 2d**). However, none of them showed enzymatic activity on native BE-activity gels, whereas all tested class I/II BEs did (**Fig. 2e**). Expression of class III BEs also did not result in glucan synthesis in yeast, as judged by the absence of dark staining after iodine staining of glucans produced in yeast cells (**Fig. 2f**). The BE activity of *At*MKIP1-YFP could not be restored by reconstituting its former catalytic triad in the *At*MKIP1(H558D,N682D)-YFP version (**Fig. 2e, f**). These data suggest that the absence of BE activity is a general feature of class III BEs and is likely due to widespread sequence divergence.

To test whether the remaining BE-like features in *At*MKIP1 are nevertheless required for its function in Arabidopsis, we mutated a glutamate and a histidine residue essential for canonical BE activity as acid/base catalyst^32^ and transition-state stabilizer^33^, respectively. Expression of this *At*MKIP1(E612A,H681A)-mCitrine construct complemented the *mkip1-1^-/-^* mutant phenotype similarly to the non-mutated *At*MKIP1-mCitrine construct (**Supplementary Fig. 2**). Complementation was also observed by *At*MKIP1(W151A,W171A)-mCitrine (**Supplementary Fig. 2**), which carries substitutions in conserved tryptophan residues critical to glucan binding of family-48 CBMs^34,35^. Taken together, these data demonstrate that *At*MKIP1 function predominantly requires others features than canonical BEs, including its newly evolved MSR domain.

### MKIP1 interacts with MatK in an RNA-independent manner

We conducted co-immunoprecipitation (IP) experiments coupled with mass spectrometry (MS) to identify potential interaction partners of *At*MKIP1. *At*MKIP1-YFP stably expressed in Arabidopsis significantly enriched *At*MatK in three independent anti-YFP IPs, each with multiple replicates (**Fig. 3a**). These experiments further suggested that *At*MKIP1 interacts with two other plastid-localized proteins essential for plant viability: *At*ValRS2 (*At*EMB2247), a putative valyl-tRNA synthetase^36^, and *At*EMB3120 (EMBRYO-DEFECTIVE 3120), a protein of unknown function^24^. We confirmed the interactions between *At*MKIP1, *At*ValRS2 and *At*EMB3120 by multiple (reverse-)IPs performed in Arabidopsis (**Fig. 3b, Supplementary Fig. 6**). However, as the functions of *At*ValRS2 and *At*EMB3120 are not or only poorly characterized, we focus here on *At*MatK.

**Fig. 3.**
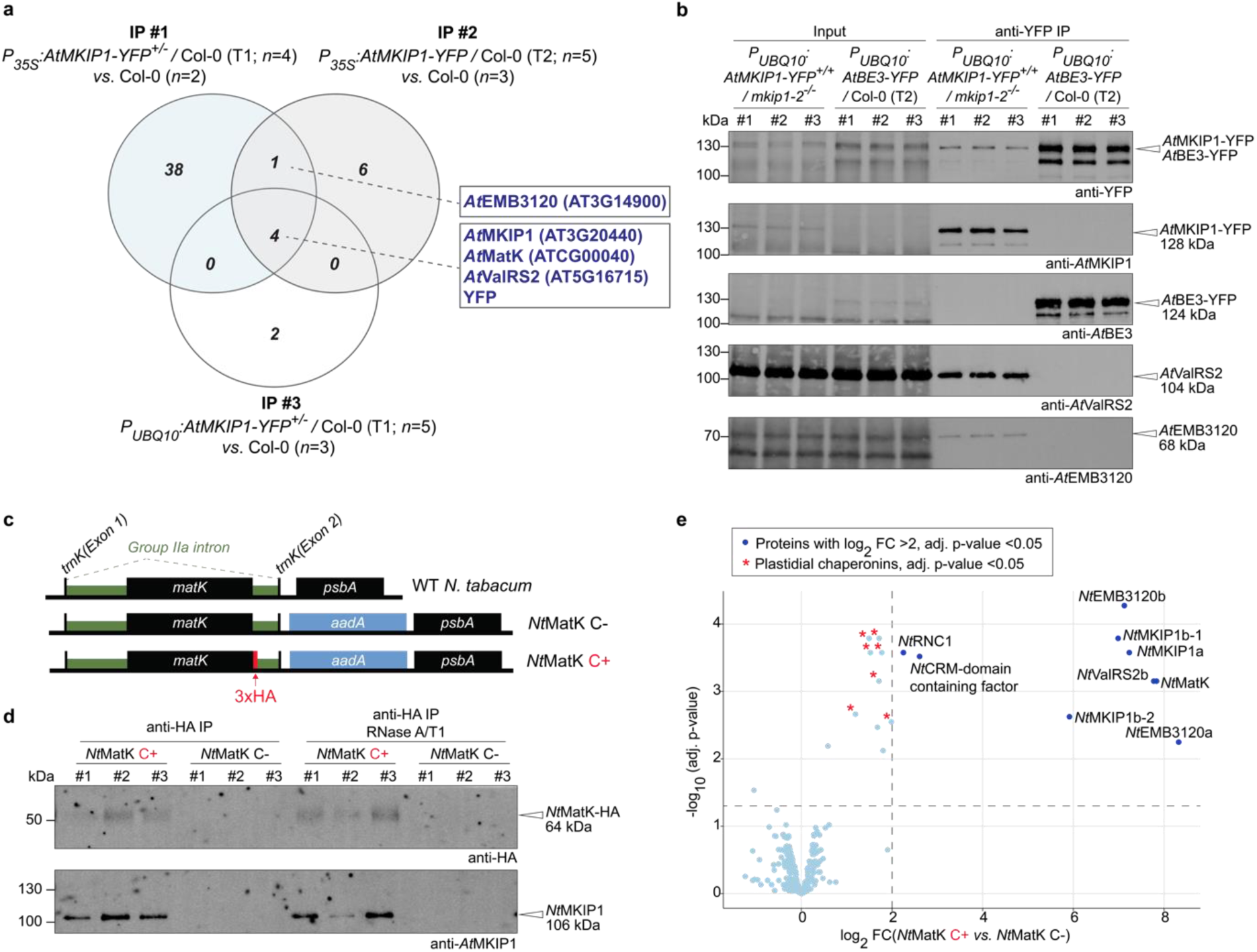
MKIP1 interacts with MatK in Arabidopsis and tobacco *in vivo*. **a** Identification of proteins associated with *At*MKIP1-YFP in mature rosettes of the indicated Arabidopsis lines. Soluble proteins were subjected to anti-YFP immunoprecipitation (IP) coupled to mass spectrometry (MS). The Venn diagram shows the number of proteins significantly enriched (log_2_ fold change ≥2, *p*-value ≤0.05) in the *At*MKIP1-YFP expressing lines compared with the wild-type (Col-0) control in three independent experiments. *n*, number of replicate plants. **b** Immunoblot detection of proteins associated with *At*MKIP1-YFP vs. *At*BE3-YFP (control) in 10-day-old Arabidopsis seedlings. Three replicates, each consisting of ∼10 g (fresh weight) of pooled 10-day old seedlings, were analyzed per line. Soluble extracts were analyzed before IP (input) and after anti-YFP IP. **c** Scheme of the *matK* plastid genome region in WT and transplastomic tobacco lines. *Nt*MatK C+ expresses *Nt*MatK with a C-terminal 3xHA tag (*red*). *Nt*MatK C- is a control line containing the *aadA* marker gene but no *matK* modification. **d** Anti-HA IP experiments using *Nt*MatK-HA as bait. Soluble protein extracted from 10-day-old *Nt*MatK C+ or *Nt*MatK C- (control) seedlings was subjected to anti-HA IP, with or without RNase A/T1 treatment during the incubation with anti-HA beads. *n*=3 seedling pools. **e** Volcano plot of proteins identified by IP-MS using *Nt*MatK-HA as bait. Anti-HA IP was performed independently from that in (**c**) but using the same workflow without RNase treatment (*n*=3 seedling pools). Proteins significantly enriched in *Nt*MatK C+ IP (adjusted p-value <0.05) with mean log_2_ fold change (FC) >2 are highlighted in dark blue. Significantly enriched proteins annotated as plastidial chaperonins are marked by red asterisks. Multiple isoforms (homoeologs) are observed because tobacco is allotetraploid. All proteins enriched in *Nt*MatK C+ IP with log_2_ FC >1 with their Arabidopsis orthologs and spectral counts are listed in **Supplementary Table 1**.

Since *At*MatK is encoded in the plastome, it cannot be routinely manipulated in Arabidopsis, and our attempts to raise a working antibody against *At*MatK were unsuccessful. We thus used *Nt*MatK C+ transplastomic tobacco (*N. tabacum*), which expresses *Nt*MatK with a C-terminal triple HA tag at its native plastome locus^11^ (**Fig. 3c**). The *Nt*MatK-HA protein is functional since it can substitute for *Nt*MatK, which is considered an essential protein^11^. In IP experiments with tobacco seedlings, *Nt*MKIP1 co-precipitated with *Nt*MatK-HA (line *Nt*MatK C+; **Fig. 3d**). This interaction was not abolished by treating the IP samples with an RNase A/T1 mixture, which effectively degraded the RNA (**Supplementary Fig. 7**), suggesting that the interaction is not bridged by (intron) RNA (**Fig. 3d**). MS analysis of an independently performed IP experiment showed a >60-fold and statistically highly significant enrichment of the bait *Nt*MatK-HA and of the tobacco orthologs of *At*MKIP1, *At*ValRS2 and *At*EMB3120 (**Fig. 3e**). In addition, *Nt*RNC1 and a CRM (chloroplast RNA splicing and ribosome maturation) domain protein, both homologs of plastidial splicing factors from maize or Arabidopsis^18,37^, were significantly, albeit weakly enriched in these IPs. Several plastidial chaperonins were also enriched (e.g., chaperonins-60 α and β; **Supplementary Table 1**), possibly indicating a need for folding assistance for *Nt*MatK-HA complex formation. However, *Nt*RNC1, the CRM domain protein and chaperonins were only enriched 2 to 6-fold, suggesting a relatively loose and/or transient association with *Nt*MatK-HA. Furthermore, none of them were enriched in the Arabidopsis IP-MS experiments (**Fig. 3a**).

We were unable to obtain high expression levels of *At*MatK in *E. coli* or from the nuclear genome *in planta* despite using a version codon-optimized for the Arabidopsis nuclear genome. However, *At*MatK with a C-terminal 3xHA tag was successfully expressed as a soluble protein in yeast, allowing us to carry out IP experiments after co-expression of *At*MatK-HA with *At*MKIP1-YFP or *At*BE3-YFP as control. *At*MKIP1-YFP as bait efficiently co-precipitated *At*MatK-HA (strain D), whereas *At*BE3-YFP did not (strain C; **Fig. 4c**). In reciprocal interaction assays, *At*MatK-HA efficiently co-precipitated *At*MKIP1-YFP, but not *At*BE3-YFP (**Supplementary Fig. 8b**). These data confirm the specific interaction between *At*MKIP1 and *At*MatK and indicate that it does not require any other plant protein or RNA.

**Fig. 4.**
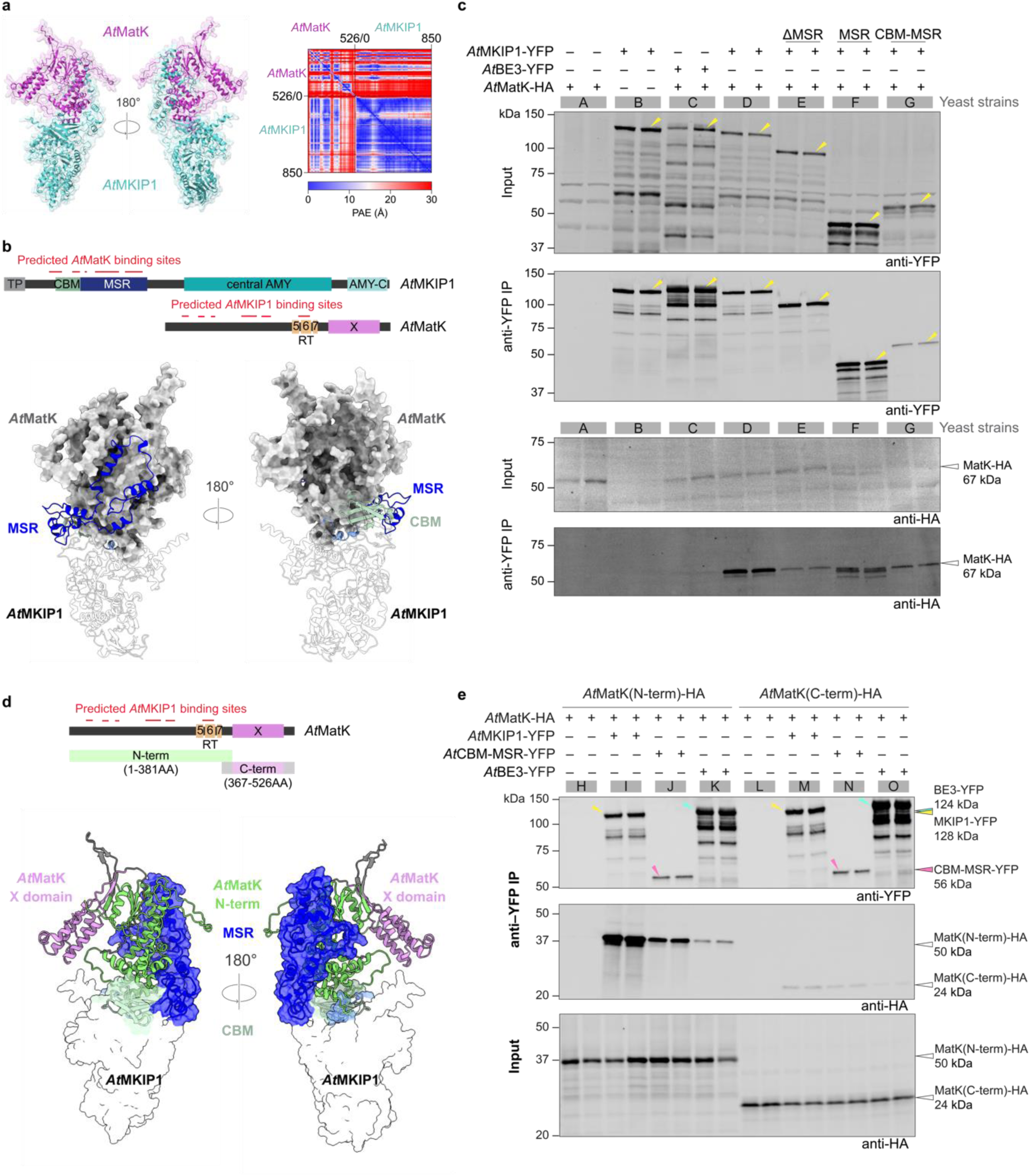
The N-terminal region of *At*MatK forms an interaction platform with the MKIP1-specific region (MSR) and carbohydrate-binding module (CBM) of *At*MKIP1. **a** AlphaFold2 prediction of the heterodimer formed between *At*MatK (*magenta*) and *At*MKIP1 (without its chloroplast transit peptide; *cyan*). The predicted aligned error (PAE) heat map (left) shows the predicted position error between all paired residues. **b** The *At*MKIP1-*At*MatK interface predicted by AlphaFold2. Top panel: Schematics of *At*MKIP1 (as in Fig. 2b) and *At*MatK. Colored boxes represent the reverse transcriptase motifs (RT5-7, *yellow*) and the X domain (*purple*). Predicted binding sites (*red*) are regions within 4 Å distance of the interacting protein. Bottom panel: Close-up view of the predicted interface, showing the protein surface of *At*MatK (*gray*) and the binding sites of *At*MKIP1 in cartoon. The binding site of *At*MKIP1 preceding the CBM is shown in light blue. **c** Immunoprecipitation (IP) of *At*MatK-HA by *At*MKIP1-YFP or *At*BE3-YFP (control) in soluble protein extracts from yeast. Arabidopsis proteins were expressed without their predicted chloroplast transit peptides, if any. The yeast background was deficient in glycogen to avoid potential interference from BEs binding to glycogen. Two replicate cultures of strains expressing the corresponding protein (+) or not (-) were analyzed. *At*MKIP1-YFP versions lacking its MSR (ΔMSR), and *At*MKIP1’s MSR and CBM-MSR (both tagged with YFP) served to test the predicted interaction platform. The soluble protein extracts were assessed before IP (input) and after anti-YFP IP using the anti-YFP or anti-HA antibodies. Yellow arrows mark the protein bands matching the molecular weights of the proteins. **d** Top panel: Schematics of *At*MatK (as in **b**). Bottom panel: Model of the interaction platform, showing the N-terminal region (*green*) and the X domain (*purple*) of *At*MatK in cartoon. **e** IP experiments performed as in (**c**) but using yeast strains expressing the N- and C-terminal *At*MatK-HA segments shown in (**d**). Anti-YFP input protein abundances are shown in **Supplementary Fig. 8c**.

AlphaFold2 multimer models predicted a heterodimer of *At*MKIP1 and *At*MatK with high confidence, as evidenced by the many intermolecular residue pairs between *At*MKIP1 and *At*MatK with low (< 5 Å) predicted aligned error (**Fig. 4a, Supplementary Fig. 9**). This model confidently predicted that the CBM-MSR region of *At*MKIP1 and the N-terminal region of *At*MatK form a large interaction platform, where ∼90 AA of both proteins engage in close (≤ 4 Å distance) intermolecular contacts (**Fig. 4b and d, Supplementary Fig. 9**). No confident heteromultimer was predicted between *At*BE3 and *At*MatK (**Supplementary** Fig. 9).

To test the involvement of these domains, we first conducted IP experiments of several *At*MKIP1 variants in yeast (**Fig. 4c)**. *At*MKIP1-YFP without MSR (ΔMSR; strain E) co-precipitated much less *At*MatK-HA than the entire *At*MKIP1-YFP (strain D). In turn, MSR-YFP co-precipitated *At*MatK-HA (strain F) only slightly less efficiently than the entire *At*MKIP1-YFP (strain D). When testing an *At*MKIP1 segment comprising its CBM and MSR, little YFP-tagged protein was expressed and recovered in the anti-YFP IP (strain G) for reasons unknown. The precipitated CBM-MSR-YFP protein, however, efficiently co-precipitated *At*MatK-HA, confirming the key role of the CBM-MSR region of *At*MKIP1 in interacting with *At*MatK.

Equivalent IP experiments using *A*tMatK-HA segments (**Fig. 4e**) showed that the N-terminal region of *A*tMatK-HA co-precipitated the whole *At*MKIP1 (strain I) and the CBM-MSR region of *At*MKIP1 (strain J) well, whereas the C-terminal X domain of *At*MatK-HA did not (strains M and N). Similarly, in reciprocal IPs using *At*MatK-HA as bait, its N-terminal portion but not the C-terminal X domain strongly co-precipitated *At*MKIP1-YFP (**Supplementary Fig. 8**). Collectively, these data suggest that the CBM-MSR of *At*MKIP1 and the N-terminal region of *At*MatK form an interaction platform, while *At*MatK’s X domain remains free, potentially to interact with intron RNA.

### *At*MKIP1 is associated with group IIA introns *in vivo*

We next conducted RNA co-IP coupled with RNA sequencing to identify RNA regions specifically associated with *At*MKIP1 or its *At*MatK complex. We therefore immunoprecipitated RNA associated with *At*MKIP1-YFP in Arabidopsis seedlings, using a line expressing *At*BE3-YFP as control. Immunoblot analysis showed the binding of *At*MKIP1’s interaction partners, confirming the success of the IP (**Fig. 3b**). Analysis of the RNA segments that were differentially enriched showed that *At*MKIP1-YFP effectively co-precipitated the RNAs of all group IIA intron containing genes previously identified to be associated with *Nt*MatK-HA^10,11^ (**Fig. 5a**). This enrichment was not due to greater abundance of these RNAs in the *At*MKIP1-YFP expressing line, as they were not or only slightly more abundant (<2-fold) in the input RNA fraction of this line. *At*MKIP1-YFP also co-precipitated RNAs of the intron-free genes *rpl23* and *rps19*, which are co-transcribed within the *rpl2*-containing operon^38^. However, while *rpl2* RNA was markedly and significantly enriched throughout the intron and exons, only short segments (< 200 nt) of *rpl23* and *rps19* flanking *rpl2* were >4-fold enriched, suggesting co-enrichment with *rpl2* (**Fig. 5b**). The lack of co-enrichment of distal transcript segments is likely due to the nuclease activities present in the chloroplast stroma and during the IP bead incubation step, which lead to RNA fragmentation except for regions protected by proteins. This enables the specific recovery of the RNA regions that are directly or indirectly (e.g., via a protein or RNA bridge) associated with the bait protein^10,11^.

**Fig. 5.**
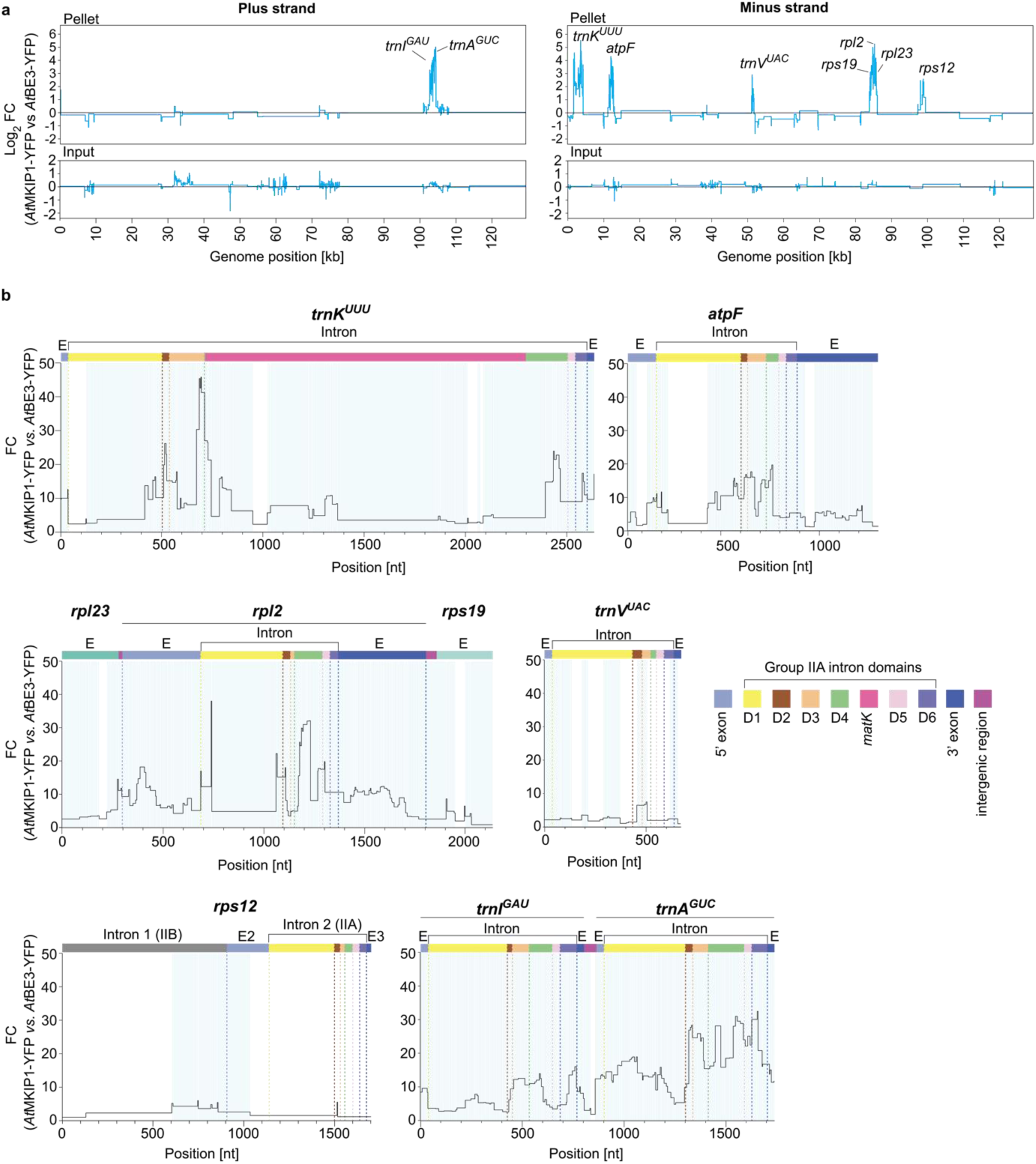
*At*MKIP1-YFP co-precipitates plastidial intron RNA. **a** Sequencing analysis of the RNA recovered by the anti-YFP immunoprecipitations (“pellets”, upper panels) shown in Fig. 3b and input RNA (lower panels). Three pools of 10-day old Arabidopsis seedlings (*P_UBQ10_:AtMKIP1-YFP* / *mkip-2* and *P_UBQ10_:AtBE3-YFP* / Col-0) were analyzed per line. Log2 fold changes (FC) indicate the mean relative abundances of the RNA segments co-precipitated by *At*MKIP1-YFP compared to those co-precipitated by *At*BE3-YFP (upper panel), or the mean relative abundance of RNA segments present in the input samples (lower panels). Relative abundances were calculated by DiffSegR, which combines RNA segments with similar fold changes into regions indicated by horizontal lines. Regions were mapped to the Arabidopsis plastome in a strand-specific manner and excluding the second copy of the inverted repeat region. Genes containing regions with a log2 FC >2 are labeled by name. **b** Zoom-in view of differentially abundant regions of the genes marked in (**a**) plotted in linear scale. Exon (E) and intron domains are annotated with color-coded boxes, with dashed lines indicating their borders. Brackets indicate group IIA introns. Significantly enriched (adjusted *p*-value <0.01) regions are highlighted with a light-blue background. Only one half of the group IIB *rps12* intron 1 is shown, as it is di-partite (trans-spliced) and the second part was not enriched.

Fine-mapping of the other RNA regions co-precipitated with *At*MKIP1-YFP showed that *trnK* enrichment peaked at the junction between intron domains III and IV, just upstream of the *matK* open-reading frame (ORF) (**Fig. 5b**). The exons and most other *trnK* intron regions were also enriched, overall largely resembling the enrichment observed by RNA co-IP of *Nt*MatK-HA^10,11^. The remaining binding targets shared a strong enrichment of intron domain II. For *rps12* intron 2, enrichment was low and domain II was the only intron domain significantly enriched, contrasting the pattern observed for *Nt*MatK-HA^10^. However, for most other targets (*atpF, rpl2, trnI*, and *trnA*), strong enrichment was also observed in intron domains IV and VI. In the case of *trnV*, enrichment was relatively low, consistent with its variable enrichment by *Nt*MatK-HA^10,11^. In addition, domain VI of *trnV* was not enriched; this domain may be minimally involved in intron splicing of *trnV* because it lacks the bulged adenosine normally used in the first splicing step^13^. Overall, these results show a remarkable overlap of the intron regions associated with *At*MKIP1-YFP and those previously identified for *Nt*MatK-HA^10,11^, suggesting a close functional link between MKIP1 and MatK.

### *At*MKIP1 facilitates the splicing of plastidial introns

To assess whether *At*MKIP1 is required for the splicing of its associated introns, we generated two estradiol-inducible artificial microRNA silencing lines in Arabidopsis, each targeting a different site of the *AtMKIP1* mRNA (*amiR-mkip1* plants). We made an equivalent line targeting *AtuL4c (AtPRPL4*) mRNA, which encodes an essential protein of the plastidial large ribosomal subunit^24^. This line served as a control for splicing defects caused by plastidial translation impairment, which could reduce *At*MatK abundance and thus the splicing efficiency of its target introns^14^.

Estradiol treatment of *amiR-uL4c* plants resulted in pale newly developed leaves, especially at the base, where the tissue is the youngest and chloroplasts are still differentiating (**Fig. 6a**). We focused on newly developed leaves (red arrows in **Fig. 6a**) in downstream analyses. This tissue was depleted of *At*uL4c protein and showed a marked reduction in plastid translation, as judged by the lower abundance of the plastome-encoded photosynthetic proteins *At*RbcL (the large Rubisco subunit) and *At*PsbA (a photosystem II subunit), as well as of the 16S and 23S rRNAs of plastidial ribosomes (**Fig. 6b-d**). Interestingly, young *amiR-uL4c* silenced leaves had higher protein levels of nuclear-encoded *At*MKIP1, *At*ValRS2 and *At*EMB3120 (**Fig. 6b**). This is consistent with their putative roles in plastid gene expression and the increased expression of such genes when chloroplast differentiation is delayed^39^.

**Fig. 6.**
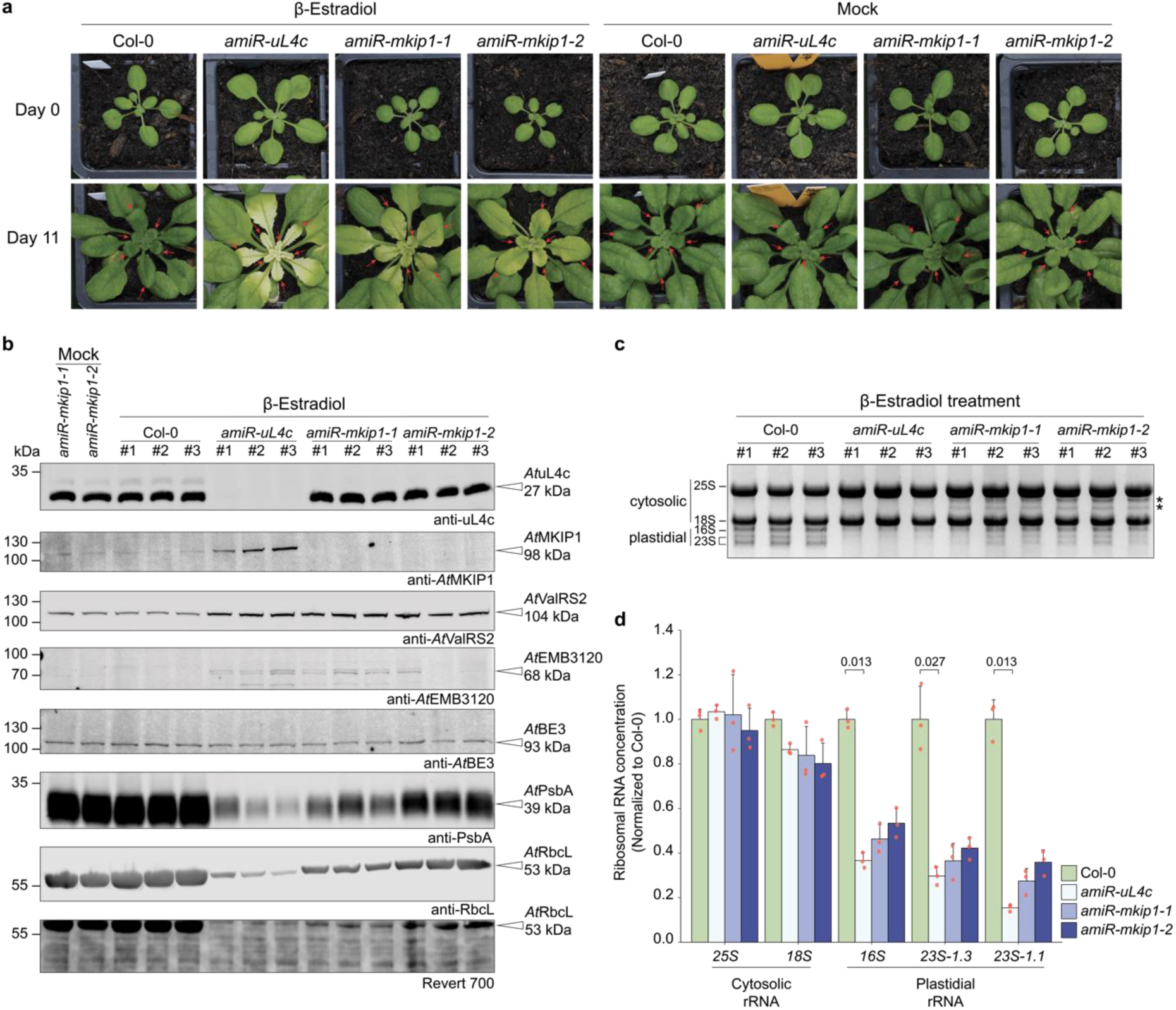
Induced silencing of *AtMKIP1* results in pale new leaves with milder plastidial translation defects than *AtuL4c* silencing. Col-0, wild type. **a** Arabidopsis rosettes before (day 0; 3-week old plants) and 11 days after daily spraying with β-estradiol or mock solution. Newly emerged leaves sampled for downstream analyses are indicated by red arrows. Replicate plants showed similar phenotypes. **b** Immunoblot analysis of protein accumulation. Total protein from newly emerged leaves after 11 days of treatment was loaded on an equal leaf area basis. Three estradiol-treated plants were analyzed per line. Revert 700, total protein stain. **c** Denaturing gel electrophoresis to assess ribosomal RNA (rRNA) accumulation. Cytosolic 25S and 18S rRNA, as well as the plastidial 16S rRNA and the 1.3 kb and 1.1 kb fragments of plastidial 23S rRNA are indicated. Equal amounts of total RNA were loaded for each sample. *n*=3 estradiol-treated plants. Asterisks mark two additional bands migrating between the 18S and 25S rRNAs (see Results for details). **d** Quantification of rRNA abundance upon β-estradiol treatment using a TapeStation device. rRNA concentrations were normalized to the mean of β-estradiol-treated wild-type samples. Shown are means ± s.d. (*n*=3 plants), red dots represent individual data points. Statistical significance was evaluated using the Kruskal-Wallis test followed by a two-sided Dunn’s post hoc multiple comparisons test and Bonferroni *p-*value adjustment. For statistically significant differences between means (*p<*0.05), the *p*-values are provided above the brackets.

In *amiR-mkip1* plants, estradiol-induced silencing reduced the abundance of *At*MKIP1, as expected (**Fig. 6b**). In addition, these plants produced pale new leaves (**Fig. 6a**) and showed a higher abundance of *At*ValRS2 and *At*EMB3120 proteins. They also showed a defect in plastid translation, but in a milder form than in the *amiR-uL4c* line, since the reduction of *At*PsbA and *At*RbcL proteins and of the plastidial rRNAs was less pronounced (**Fig. 6b-d**). In silenced *amiR-mkip1* tissue, we additionally observed two rRNA bands migrating between the 25S and 18S rRNA (**Fig. 6c**), with sizes corresponding to plastidial 23S rRNA intermediates likely resulting from incomplete “hidden breaks”, indicating impaired ribosome assembly^40^. As for plastid translation defects, the alteration of rRNA processing does not necessarily indicate a direct role in this process but is a frequent secondary effect of diverse defects in plastid gene expression^1^. In silenced *amiR-uL4c* tissue, these intermediates did not accumulate, presumably due to the greater overall reduction in plastid rRNAs (**Fig. 6d**).

We next assessed the efficiency of intron splicing in newly emerged, silenced tissue. We therefore quantified the relative ratios of spliced to unspliced plastid mRNAs using reverse transcription quantitative real-time PCR (RT-qPCR)^41,42^. Given the difficulties associated with reverse transcription of tRNAs, their splicing was determined by northern blotting, circumventing cDNA synthesis. Splicing of the group IIB intron in *rps16* was not assessed, since this likely is a pseudogene in Arabidopsis with no apparent splicing even in the WT^43^.

No significant splicing defects compared to WT were observed in mock-treated plants of any *amiR* lines (**Fig. 7b**, **Supplementary** Figs. 10**, 11)**. However, upon estradiol-induced silencing, young *amiMKIP1* tissue showed major splicing defects in most of the introns identified to be associated with *At*MKIP1-YFP (**Fig. 5**). The introns of *trnK*, *trnA*, *rps12* (intron 2), *rpl2*, and *atpF* showed severely reduced splicing efficiency (i.e., a lower ratio of spliced to unspliced RNA) and less absolute spliced RNA compared to estradiol-treated WT and *amiR-uL4c* plants (**Fig. 7, Supplementary Fig. 10**). Mild splicing defects in *AtMKIP1* silenced tissue were also observed for *trnI* and *trnV* (**Fig. 7b, c**). In contrast, none of these MatK binding targets showed strongly impaired splicing in *AtuL4c* silenced tissue (**Fig. 7, Supplementary** Fig. 10), indicating that *At*MatK abundance was still sufficient. This is in agreement with a previous report of Arabidopsis that MatK targets are correctly spliced despite impaired plastid translation^26^.

**Fig. 7.**
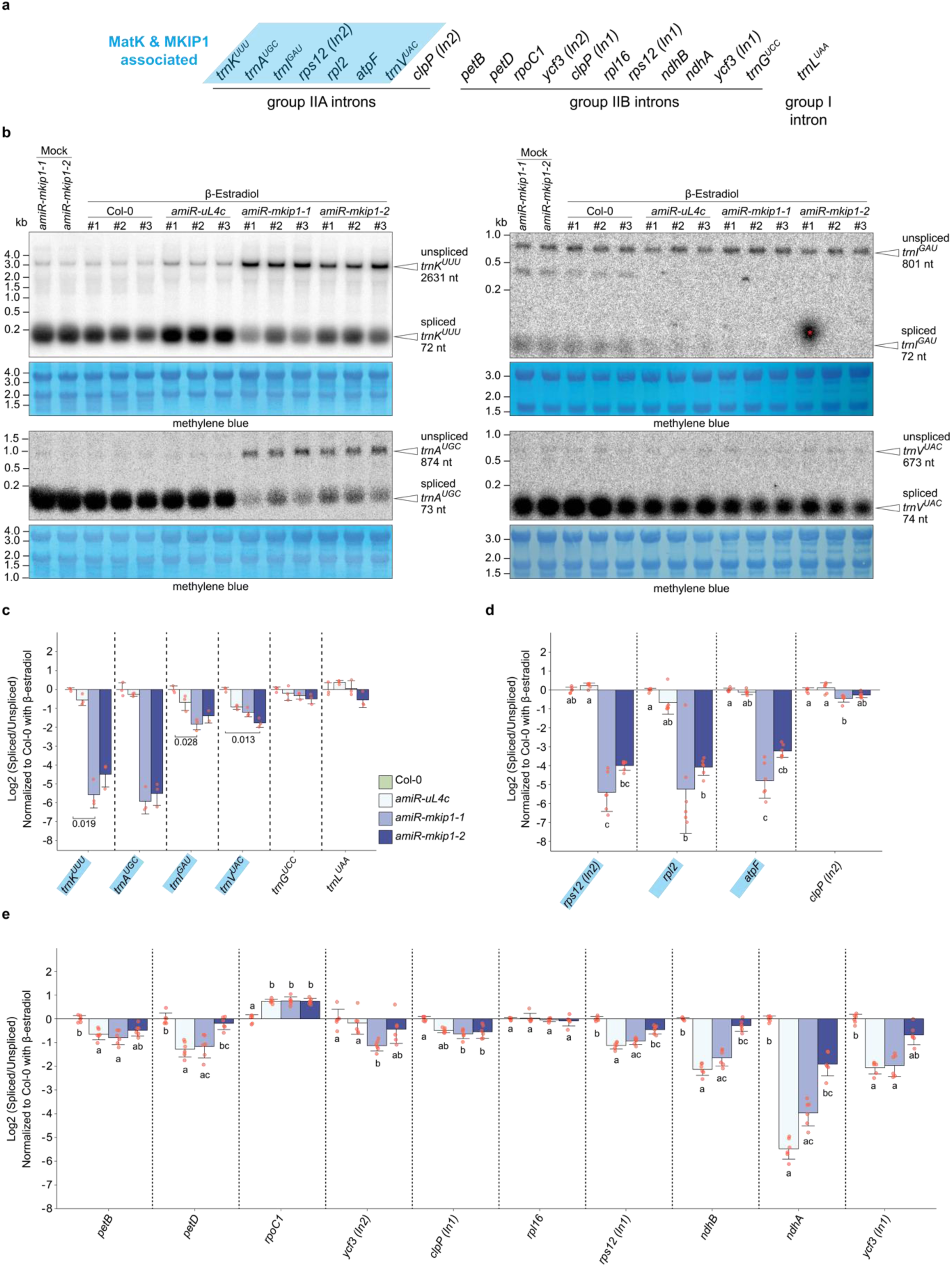
Silencing of *AtMKIP1* leads to splicing defects in MKIP1/MatK-associated introns. β-estradiol treatment and sampling of newly emerged leaves were performed as described for Fig. 6. Col-0, wild type. **a** Scheme of the introns present in the chloroplast genome. MKIP1/MatK-associated introns are highlighted in blue. **b** Northern blot analysis of intron splicing in tRNAs using probes against exons 1 of *trnK^UUU^*, *trnA^UGC^*, *trnI^GAU^*, or *trnV^UAC^*. Methylene blue is a total RNA stain used as a loading control. **c** Intron splicing efficiency of tRNAs in estradiol-treated plants. MKIP1/MatK associated RNAs are highlighted in blue. Efficiencies are the log2 transformed ratios of spliced to unspliced transcripts of a given sample normalized to that of the estradiol-treated Col-0 mean, based on the data presented in (**b**) and **Supplementary** Fig. 11. Shown are means ± s.d. (*n*=3 plants), except for *trnI*, where one replicate with non-specific contamination (marked with a red asterisk in **b**) was excluded. Red dots indicate single data points. Statistical significance was evaluated using the Kruskal-Wallis test followed by a two-sided Dunn’s post hoc multiple comparisons test and Bonferroni *p* value adjustment. For statistically significant differences between means (*p<*0.05), the *p*-values are provided above the brackets. **d-e** Intron splicing efficiency of mRNAs in estradiol-treated plants assessed by RT-qPCR. Splicing efficiency and statistical significance were calculated as in (**c**). Shown are means ± s.d. (*n*=6 plants). Different letters indicate statistically significant differences (*p<*0.05) between the means of the β-estradiol-treated samples. Underlying abundances of spliced and unspliced mRNAs are presented in **Supplementary** Fig. 10.

The ratio of spliced to unspliced RNA upon *AtMKIP1* silencing was also somewhat reduced in several introns not associated to *At*MKIP1-YFP (**Fig. 7e**). However, in all cases other than the *petB*, *petD* and *ndhA* intron, this was due to the accumulation of the unspliced precursor, while the spliced RNA appeared not to be reduced compared to WT (**Supplementary Fig. 10**). In addition, here *AtuL4c* silencing caused analogous defects, suggesting that these are secondary effects associated with chloroplast malfunction^2^. Collectively, these findings strongly suggest that *AtMKIP1* silencing causes a specific and primary splicing defect in all introns associated to MatK and MKIP1. Given that these include tRNAs and transcripts encoding ribosomal proteins, the splicing defects likely underlie the observed plastidial translation defects, ultimately leading to pale leaves in the *amiR-mkip1* plants, and, in the *mkip1^-/-^* lines, to the early embryonic defects.

## Discussion

*At*MKIP1 has been suggested to function in plastid carbohydrate metabolism, largely based on its annotation as a BE and the reduced starch content of a hypomorphic *mkip1* mutant^23,44^. The observation that MKIP1 proteins have lost catalytic activity on glucans^21,22^ (**Fig. 2**) renders such an involvement unlikely. Instead, our work provides several lines of evidence that the essential function of *At*MKIP1 is to assist plastidial splicing in conjunction with *At*MatK. First, MKIP1 physically interacts with MatK, not requiring a protein or RNA bridge (**Figs. 3** and **4**). Second, RNA-IP experiments demonstrate that *At*MKIP1 is associated with the same intron RNAs as *Nt*MatK^10,11^ *in vivo* (**Fig. 5**). Third, induced *AtMKIP1* knockdown results in a primary splicing defect specifically in all MatK binding targets (**Fig. 7**), reinforcing a direct role of *At*MKIP1 in the splicing of *A*tMatK targeted introns.

A function in plastid intron splicing helps to explain the drastic consequences on plant viability observed upon *At*MKIP1 loss, since defects in plastid gene expression in Arabidopsis lead to embryonic defects and frequently lethality^36,37^. It is also consistent with the apparent emergence of the *MKIP1* gene within streptophyte algae (**Fig. 2**), which coincides with the acquisition of the plastidial *matK* gene and group IIA introns^12^. In agreement with this, we identified at least one *MKIP1* gene in all land plant species assessed, except for two non-photoautotrophic parasitic *Cuscuta* species that also lost the *matK* gene and its associated plastid introns^45^, possibly rendering MKIP1 dispensable (**Supplementary Fig. 3a**). In contrast to canonical BEs, MKIP1 has also not been identified in starch-associated proteomes^46,47^ but instead in the “core” set of ∼200 plastidial nucleoid-associated proteins in maize and Arabidopsis^48,49^, together with MatK and other proteins involved in gene expression. *At*MKIP1-YFP did not show the punctate suborganellar distribution pattern typical of nucleoids (**Fig. 1b**). However, the low signal of *P_UBQ10_::At*MKIP1-YFP necessitated strong *P_35S_*-driven overexpression, which could have caused (partial) mislocalization if nucleoid association requires complex formation with *At*MatK or other proteins that are lower in abundance.

The neofunctionalization of MKIP1 likely was a multi-step adaptation process. A suggestive feature is its MSR that protrudes from the *At*MKIP1 core. The origin of the MSR is unclear, as we could not find any homologous protein or DNA sequences beyond class III BEs. Nevertheless, it is essential for the function of *At*MKIP1 **(Supplementary** Fig. 5), probably because it confers – together with the CBM – the interaction with *At*MatK (**Fig. 4**). The MSR may also interact with RNA; it has conserved lysine and arginine residues at its C-terminal end (**Supplementary** Fig. 4), which are predicted to be surface exposed in the dimer. Interestingly, the CBM appears to have adapted to MKIP1’s new role. It is unlikely to function in glucan binding, since mutations in the tryptophan residues normally involved in carbohydrate binding to alanine did not impact the function of *At*MKIP1 (**Supplementary Fig. 2**). Rather, in MKIP1 proteins, the CBM has been repurposed to become part of the MatK-interaction platform, and the *At*MKIP1-*At*MatK dimer model predicts that the CBM is now oriented toward *At*MatK, forming part of the extensive protein-protein interface (**Fig. 4**). If correct, this would preclude the aforementioned tryptophans from making contact with glucans. MKIP1 proteins also display extensive sequence divergence across the central and C-terminal α-amylase domains and have lost the integrity of the former catalytic center, rendering them enzymatically inactive as a BE (**Fig. 2**). These changes may have contributed to the elimination of glucan interaction sites on the protein surface^50^, further diverging the function of MKIP1 from that of canonical BEs. However, the α-amylase domains are likely to support MKIP1 functions in other, as yet unknown ways, as they are under evolutionary constraint and a P749S substitution in the central α-amylase domain of *At*MKIP1 resulted in pale leaves^44^.

MKIP1 is analogous to other enzyme(-like) factors recruited to facilitate plastidial intron splicing, but which no longer require or possess enzymatic activity, including the pseudouridine synthase homolog Raa2, implicated in group IIA trans-splicing in *Chlamydomonas*^51^, and maize CHLOROPLAST RNA SPLICING2 (CRS2), a catalytically-dead peptidyl-tRNA hydrolase required for group IIB intron splicing^52^. There are also several nuclear-encoded factors that assist in the splicing of MatK-associated group IIA introns^1,2^, some of which possess protein domains originally associated with other functions (e.g., RNC1 and CRS1 from maize^17,18^). However, none of these factors were reported to be in direct contact with MatK, consistent with their weak or absent enrichment in the *Nt*MatK-HA IP (**Fig. 3e**), nor do they associate with the very same introns as MatK. This is consistent with upstream roles in intron folding and/or RNA stabilization.

By contrast, the physical and functional association of MKIP1 with MatK suggests a role of MKIP1 in directly helping MatK function (**Fig. 8**). The sequence divergence of MatK’s N-terminal region away from its ancestral RT domain implies that it cannot fulfill its original function in establishing the intron-specific contacts^6,7,15^. Strikingly, our data provides strong evidence that this region now interacts with MKIP1. Complex formation may functionally compensate for the loss of MatK’s N-terminal domain, thereby helping its initial interaction with RNA and/or with intron folding into a catalytically competent structure. The complex may equip MatK with a larger area for RNA binding, which may enable it dock to multiple introns via less specific contacts, and thus help it to act as a general splicing factor. In bacterial maturases, the high-affinity binding of the RT domain to intron domain IV also serves to occupy the ribosome binding site and the start codon of the maturase ORF, so that translation does not interfere with the splicing process^53^. Among the MatK-associated introns, the *trnK* intron has retained an ORF, whose start and immediate upstream region were strongly associated with *Nt*MatK^10^ and *At*MKIP1 (**Fig. 5b**), which might enable MatK to regulate its own translation. MatK autoregulation is supported by mathematical models of MatK expression^54^ but was not observed upon co-expression of MatK with *trnK* intron regions in *E. coli*^55^, potentially due to a requirement of MKIP1 for binding to this site. At the same time, our data also imply that the X domain of MatK remains free of MKIP1 interactions, allowing it to fit into the intron core. Given the high degree of structural conservation of the domain X in MatK^15^, it is likely to function analogously to bacterial maturases in facilitating conformational RNA changes during splicing^9^.

**Fig. 8.**
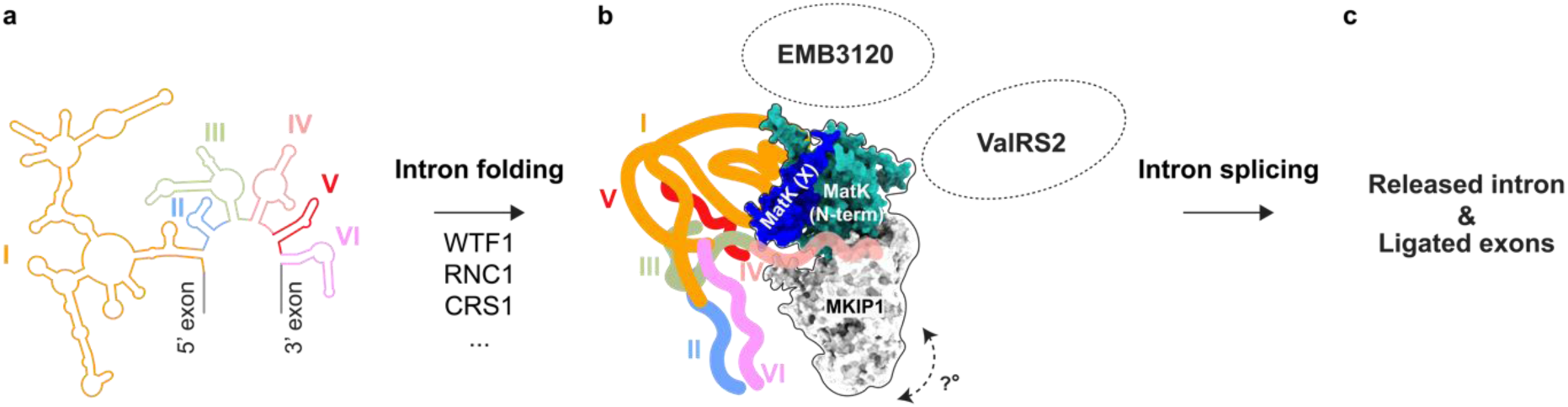
Proposed role of the MatK complex during plastidial group IIA intron splicing. **a** Variable sets of nuclear-encoded proteins (such as CRS1 and/or WTF1 and RNC1, refs. 17-19) facilitate initial intron folding and/or stabilize the intron (exemplified by the Arabidopsis *atpF* intron, with intron domains indicated by numbers). **b** MatK, MKIP1 and possibly also EMB3120 and ValRS2 bind to the pre-folded intron to promote the RNA-catalyzed intron splicing as well as further intron folding and/or stabilization. Initial RNA binding likely involves interaction between the MatK complex and intron domain IV (*salmon*), allowing the X domain of MatK (*dark blue*) to make contact with the catalytic core of the intron RNA and to promote intron splicing. The modelled positions of the RNA, MatK and MKIP1 are based on the cryo-EM structure of a bacterial group IIA intron–maturase complex^8^ and the AlphaFold2 prediction of the MatK-MKIP1 dimer. The angle of MKIP1 (*grey*) and the N-terminal domain of MatK (N-term, *cyan*) relative to the X domain of MatK and the RNA could not be confidently predicted. Exons are not shown. **c** Splicing results in the release of the intron, the spliced transcript and the MatK complex (not shown).

RNA binding and splicing may also be assisted by additional interaction partners. Our observation that ValRS2, EMB3120, MKIP1 and MatK were all highly enriched in the tobacco and Arabidopsis IPs (**Fig. 3**) suggests that they altogether form a complex. If ValRS2 does indeed play a role in MatK-mediated splicing, it probably has retained its function as a valyl-tRNA synthetase, as it is the only plastid-localized protein with such predicted function^36^. Its recruitment as a splicing factor would thus resemble that of bifunctional tRNA synthetases in *Neurospora*^56^ and yeast^57^, which help the splicing mitochondrial group I introns in addition to having tRNA synthetase function. While the intron associations of *At*MKIP1 and splicing impairments upon *AtMKIP1* silencing suggest that MKIP1 is part of the MatK complex for all of its targets, EMB3120 and ValRS2 might only associate during the splicing of some introns. Alternatively, the composition of the MatK splicing complex could vary during the sequential steps of splicing. The latter situation would resemble the major type of nuclear spliceosomes, which are also thought to derive from group II introns^3^; these have the same composition regardless of the intron, but undergo complex remodeling during each splicing step. Further work will be required to dissect the (dynamic) contribution of each component of the MatK complex during intron binding, folding and splicing.

## Methods

### Mutant lines and generation of transgenic plants

Unless indicated otherwise in the figures, experiments were performed on Arabidopsis single-insertion lines of the T3 or a later generation. Where zygosity is not indicated in the Figures, plants were either homo- or heterozygous for the insert. Binary plant expression constructs were transformed into *Agrobacterium tumefaciens* (strain GV3101). *Arabidopsis thaliana* (Col-0) plants were transformed by floral dipping. Transformed T1 plants were selected for their resistance to hygromycin or Basta on 0.8% (w/v) agar plates containing one-half-strength Murashige and Skoog (MS) medium including vitamins (Duchefa Biochemie) at pH 5.7 with 25 mg/L hygromycin B or 15 mg/L Basta and further tested for transgene expression by immunoblotting. Single insertion lines were identified based on the segregation ratio of resistant to non-resistant plants in the T2 generation.

T-DNA insertion lines were in Col-0 background from the SALK collection and were obtained from the European Arabidopsis Stock Centre under the following identifiers: SALKseq_040659 (*mkip1-1*), SALK_129083 (*mkip1-2* or *emb2729-2*) and SALK_070161 (*valrs2-1*). Primers used for genotyping are listed in **Supplementary Data 1**. Insertion sites were confirmed by Sanger sequencing of the PCR amplicon containing the border between the T-DNA and its genomic insertion site.

Homoplastomic *Nicotiana tabacum* (tobacco) lines modified to express *Nt*MatK fused to a C-terminal 3x HA tag (line *Nt*MatK C+) or control lines carrying only the selection marker (*Nt*MatK C-)^11^ were kindly provided by Christian Schmitz-Linneweber (HU Berlin, Germany).

### Cloning of plant constructs

Unless otherwise noted, all plasmids for protein expression in Arabidopsis were generated using Gateway recombination cloning technology (Invitrogen). The plasmids were validated by sequencing of the insert(s) (when PCR-amplified fragments were cloned) and/or diagnostic restriction digests (when PCR was not used).

The CDS of the full-length primary splice forms of *AtMKIP1* (AT3G20440.2), *AtEMB3120* (AT3G14900.1), and *AtValRS2* (AT5G16715.1) flanked by attB sites were amplified by PCR from Arabidopsis Col-0 cDNA and individually recombined into pDONR221. The resulting vectors (AtMKIP1_pDONR221, AtEMB3120_pDONR221, and AtValRS2_pDONR221) were separately recombined with pUBC-YFP^58^, which allows *P_UBQ10_*-driven expression of proteins carrying a C-terminal YFP tag. For P*_35S_*-driven expression of *At*MKIP1 with a C-terminal YFP tag, AtMKIP1_pDONR221 was recombined with the gateway vector pB7WG2.

For expression of *At*MKIP1 with a C-terminal mCitrine tag driven by *P_ABI1_* or *P_UBQ10_*, we first amplified the respective promoter sequences (for *P_ABI1_*: 2.2 kb preceding the start codon of *ABI3,* AT3G24650; for *P_UBQ10_*: the 0.6 kb promoter as in pUBC-YFP) flanked by attB sites by PCR using Arabidopsis Col-0 genomic DNA or the pUBC-YFP vector as templates. After recombining the PCR products into pDONR P4-P1R, the resulting promoter vectors (*PABI3_*pDONRP4-P1R and PUBQ10_pDONRP4-P1R) were separately recombined with mCitrine_pDONR P2R-P3 (ref. ^35^) and AtMKIP1_pDONR221 into the gateway multi-site destination vector pB7m34GW2.

Point mutations in the CBM of *At*MKIP1 (W151A, W171A) or the central α-amylase domain of *At*MKIP1 (E612A, H681A) were introduced into AtMKIP1_pDONR221 using two sequential rounds of site-directed mutagenesis for each substitution (QuikChange; Agilent Technologies). The resulting AtMKIP1(W151A,W171A)_pDONR221 and AtMKIP1(E612A,H681A)_pDONR221 vectors were then separately recombined with PUBQ10_pDONRP4-P1R and mCitrine_pDONR P2R-P3 (ref. ^35^) into pB7m34GW2. To express a ΔMSR version of *At*MKIP1, the MSR (AA 178-333 of *At*MKIP1) in AtMKIP1_pUBC-YFP was deleted using NEBuilder HiFi DNA Assembly (New England Biolabs) by fusing PCR-amplified vector fragments covering the entire vector except for the MSR, resulting in pBP605_AtMKIP1_deltaMSR(178-333)_pUBC-YFP.

amiRNAs (21mers; **Supplementary Data 1**) for inducible silencing of *AtMKIP1* or *AtuL4c* were designed using the WDM3 MicroRNA Designer tool. Synthesized amiRs in the *MIR319a* backbone flanked by attB sites were first recombined into pDONR221 and then into binary pH7m34GW vectors^59^ containing an estrogen-inducible XVE cassette driven by *P_UBQ10_*.

### Plant growth and β-estradiol treatment

Arabidopsis and tobacco plants were grown in Percival AR-95L chambers (CLF Plant Climatics) under 12-h light and 12-h dark cycles as previously described^60^. For β-estradiol-induced silencing, plants were sprayed once per day with aqueous β-estradiol solution (20 µM β-estradiol [Sigma-Aldrich], 0.2% [v/v] dimethyl sulfoxide [DMSO], 0.01% [v/v] Silwet L-77). The mock spray solution was prepared with the same composition but omitting β-estradiol.

### Plant imaging

For imaging Arabidopsis chloroplasts by transmission electron microscopy (TEM), Arabidopsis leaf sections were sampled from 3-week-old soil-grown plants. Leaf sections were fixed, embedded, and imaged as described previously^60^. For observing *At*MKIP1-YFP localization, leaves of 3-week-old soil-grown Arabidopsis plants were imaged using a Zeiss LSM780 confocal microscope and a 40× water objective using the 514-nm (for YFP) and the 633-nm (for chlorophyll autofluorescence) helium-neon lasers as light source.

To image Arabidopsis embryos, seeds dissected from a silique were collected in a microcentrifuge tube and incubated in a 1:1 mixture of ethanol and acetic acid for 1 h at room temperature with rotation, then placed on a microscope glass slide and further cleared by incubation with Hoyer’s medium (7.5 g gum arabic, 100 g chloral hydrate, 5 ml glycerol and 30 ml water) for 1 h. Cleared seeds were imaged using a Leica DM2500 microscope.

### Plant protein extraction and immunoblotting

Total plant protein extraction from leaf disks and immunoblotting using IRDye 800CW goat anti-rabbit IgG secondary antibody (LICORbio) were conducted as described previously^60^. Total proteins on membranes were stained with Revert 700 (LICORbio). Antibody sources, dilutions and validation are detailed in **Supplementary Data 1**.

### Immunoprecipitation in plants for immunoblot detection

For anti-YFP IP experiments in Arabidopsis, freshly harvested plant tissue was homogenized in ice-cold extraction buffer (50 mM Tris–HCl, pH 8.0, 150 mM NaCl, 1 mM DTT, 1% [v/v] Triton X-100, 1x protease inhibitor cocktail [cOmplete EDTA-free, Roche]) using equal ratios of buffer to input material for all samples and a Dounce homogenizer. After centrifugation at 19,000 *g* and 4°C for 5 min, the supernatant containing soluble protein was incubated with pre-equilibrated magnetic agarose anti-YFP beads (GFP-Trap, ChromoTek; 10 µl beads per 0.1 g plant material) for 1 h at 4°C with rotation, washed three times with ice-cold wash buffer 1 (same as extraction buffer, but replacing Triton X-100 with 0.5 % [v/v] NP-40), followed by three washes with ice-cold wash buffer 2 (50 mM Tris–HCl, pH 8.0, 150 mM NaCl), each time collecting the beads on a magnetic rack. In the last wash step, the resuspended beads were transferred to a fresh tube to remove contaminants sticking to the tube. Proteins bound to the beads after washing were eluted by boiling the beads in SDS-PAGE loading dye (50 mM Tris-HCl, pH 6.8, 2% [w/v] SDS, 100 mM DTT, 3% [v/v] glycerol, and 0.005% [w/v] bromophenol blue) at 95°C for 5 min. Input samples are aliquots of soluble proteins not subjected to immunoprecipitation.

For the anti-HA IP in *N. tabacum*, 10-day-old seedlings were used as the starting material. Soluble protein extraction and bead incubation were carried out as described above for Arabidopsis samples but using anti-HA beads (Anti-HA high affinity, Roche). After protein binding during the incubation step, the beads were collected by centrifugation at 13,000 *g* for 10 s, then washed three times with extraction buffer and three times with dilution buffer (50 mM Tris–HCl, pH 8.0, 150 mM NaCl). In the final wash step, the resuspended beads were transferred to a new tube and collected by centrifugation. Bound proteins were eluted as for Arabidopsis. If samples were RNase treated, RNase A/T1 mix (Thermo Fisher) was added to the soluble protein extracts (final concentrations: 100 µg/ml for RNase A and 250 U/ml for RNase T1) together with the anti-HA beads.

### Immunoprecipitation in plants followed by mass spectrometry

For IP-MS experiments in Arabidopsis, we followed a procedure previously described^35^ using anti-YFP µMACS magnetic beads (Miltenyi Biotec) and protein identification on a Q Exactive mass spectrometer (Thermo Scientific) with the following modifications: 1) proteins eluted from the beads were precipitated in 20% (w/v) trichloroacetic acid; 2) when needed, peptides were cleaned using the Phoenix kit (PreOmics) following the manufacturer’s instructions; 3) dried peptides were solubilized in 3% (v/v) acetonitrile, 0.1% (v/v) formic acid prior to MS analysis; 4) liquid chromatography of peptide separation was conducted at a flow rate of 300 nl/min by a gradient from 5% solvent B (0.1% formic acid in acetonitrile)/95% solvent A (0.1% formic acid in water) to 35% B / 65% s A in 90 min, to 60% B/ 40% A in 5 min and to 80% B/20% A in 1 min.

The acquired raw MS data were processed by MaxQuant (v1.6.2.3), followed by protein identification using the integrated Andromeda search engine. A separate MaxQuant analysis was performed for each of the three IP experiments. Spectra were searched against the Araport 11 database (v2017-02-01), concatenated to its reversed decoyed FASTA database and common protein contaminants, setting cysteine carbamidomethylation as fixed and methionine oxidation and N-terminal protein acetylation as variable modifications, enzyme specificity to trypsin and allowing a maximum of two missed-cleavages, and using MaxQuant Orbitrap default search settings. The maximum false discovery rate was set to 0.01 for peptides and 0.05 for proteins, label free quantification enabled and a 2-minutes window for matching between runs applied. Each file was kept separate in the MaxQuant experimental design to obtain individual quantitative values. Fold changes of protein abundances were computed based on intensity values reported in the proteinGroups.txt file. A set of functions implemented in the R package SRMService was used to filter for proteins with 2 or more peptides allowing for a maximum of up to 3 missing values, to normalize the data with a modified robust z-score transformation and to compute *p*-values using t-tests with pooled variance. If all measurements of a protein were missing in one of the conditions, a pseudo fold change was computed replacing the missing group average by the mean of 10% smallest protein intensities in that condition.

For IP experiments in tobacco followed by MS analysis, anti-HA IP on tobacco seedlings using anti-HA beads (Anti-HA high affinity, Roche) was conducted as for immunoblot detection (described above) without RNase treatment. After the final wash, the beads were re-suspended in 45 µl digestion buffer (triethylammonium bicarbonate, pH 8.2), reduced with 5 mM tris(2-carboxyethyl)phosphine and alkylated with 15 mM chloroacetamide. Proteins were on-bead digested overnight at 37°C using 5 µl of sequencing-grade trypsin (100 ng/µl in 10 mM HCl, Promega). The supernatants were transferred into new tubes, the beads washed with 150 µl trifluoroacetic acid (TFA) buffer (0.1% TFA, 50% acetonitrile) and combined with the first supernatant. After drying, the samples were re-solubilized in 3% (v/v) acetonitrile, 0.1% (v/v) formic acid and analyzed on an Orbitrap Exploris 480 mass spectrometer (Thermo Fisher Scientific) equipped with a Nanospray Flex Ion Source (Thermo Fisher Scientific) and coupled to an M-Class UPLC (Waters). Peptides were loaded on a nanoEase MZ Symmetry C18 trap column (100 Å, 5 µm, 180 µm x 20 mm, Waters) followed by a nanoEase MZ C18 HSS T3 column (100 Å, 1.8 µm, 75 µm x 250 mm, Waters). Peptides were separated at a flow rate of 300 nl/min by the following gradient: initial hold at 5% solvent B (0.1% formic acid in acetonitrile)/95% solvent A (0.1% formic acid in water) for 3 min, followed by a gradient to 22% B / 65% A in 43 min, and to 32% B / 68% A in 5 min, a cleaning step at 95% B for 10 min and re-equilibration at 5% B / 95 % A for 10 min. MS settings were as follows: data-dependent mode (DDA) with a maximum cycle time of 3 s, acquisition of full-scan MS spectra (350−1,200 m/z) at a resolution of 120,000 at 200 m/z after accumulation to a target value of 3,000,000 or for a maximum injection time of 50 ms, selection of precursors with an intensity > 5,000 for MS/MS. Ions were isolated with a 1.2 m/z isolation window, fragmented by higher-energy collisional dissociation using a normalized collision energy of 30%, spectra acquired at 30,000 resolution, with maximum injection time set to auto. Singly, unassigned and charge states higher than six were rejected. Precursor masses previously selected for MS/MS measurement were excluded from further selection for 20 s, and the exclusion window was set at 10 ppm. The samples were acquired using internal lock mass calibration on m/z 371.1012 and 445.1200.

The mass spectrometry data were further processed using Philosopher followed by label-free quantification based on precursor intensities using IonQuant. MS2 spectra were searched against the *N. tabacum* UniProt reference proteome (UP000084051; concatenated to its reversed decoyed database and supplemented with MKIP1 proteins annotated in *N. sylvestris* [A0A1U7XZG9_NICSY, A0A1U7Y1B8_NICSY] and common contaminants by the MSFragger search engine (v3.4), allowing for one missed tryptic cleavage and fixed carbamidomethylation of cysteine, variable methionine oxidation, and variable acetylation of the protein N-terminus after methionine removal. Quantification was performed by IonQuant (v1.7.17), applying the false-discovery-rate controlled match-between-runs algorithm. Fold changes and adjusted *p*-values were calculated using the Amica webserver (v3.0.1).

### RNA co-immunoprecipitation and sequencing

Soluble cell components were extracted by homogenizing 10 g of seedlings per replicate in a pre-cooled blender using 30 ml of ice-cold extraction buffer (50 mM Tris–HCl, pH 8.0, 150 mM NaCl, 1 mM DTT, 1% (v/v) Triton X-100, 1 mM EDTA, 4 mM MgCl_2_, 1x protease inhibitor cocktail [cOmplete EDTA-free, Roche], 40 U/ml RNase inhibitor [RiboLock RNase Inhibitor, Thermo Fisher Scientific]). The homogenates were filtered through two layers of Miracloth (Milipore) and cleared by two sequential centrifugation rounds at 4,000 *g* for 10 minutes at 4 °C. Per replicate, 20 ml of the supernatant were incubated with 150 µl pre-equilibrated magnetic agarose anti-YFP beads (GFP-Trap, ChromoTek) for two hours at 4 °C with rotation, washed three times with ice-cold wash buffer 1 (same as extraction buffer, but replacing Triton X-100 with 0.5 % [v/v] NP-40), followed by three washes with ice-cold wash buffer 2 (50 mM Tris–HCl, pH 8.0, 150 mM NaCl). In the last wash, 10% (v/v) of the resuspended beads were collected in a separate tube and used for immunoblot analysis, while the residual beads were transferred to a new microcentrifuge tube and were processed for RNA extraction. For RNA extraction, 500 µl of input soluble protein extract or IP-processed beads were brought to a total volume of 1 ml with DEPC-treated water, mixed with 1 ml phenol:chloroform:isoamyl alcohol (Roth) and subjected to phenol-chloroform RNA extraction as described previously^61^. Two µg input RNA was treated with TURBO DNase (Thermo Fisher Scientific) in a 50 µl reaction according to the manufacturer’s instructions. From this, 600 ng purified RNA was subjected to rRNA depletion treatment as previously described^62^. The RNA was eluted in 18 µl of RNase-free water and supplemented with 2 µl RNA fragmentation buffer (400mM Tris-acetate pH 8.3, 1 M potassium acetate, 300mM magnesium acetate), followed by incubation at 94 °C for 5 minutes. Both pellet and input RNA were treated with T4 polynucleotide kinase (PNK; Thermo Fisher Scientific) as previously described^62^.

Library preparation for RNA sequencing was performed using the NEXTflex small RNA-seq kit v3 (Perkin Elmer) according to the manufacturer’s instructions. The resulting cDNA was amplified with a barcode incorporated in the primer and purified according to the instructions of the NEXTflex kit. Libraries were pooled for single-end 100-bp sequencing in a NovaSeq 6000 machine. Sequencing reads were adapter-trimmed and filtered for size between 18 and 60 nt using Cutadapt. The 8-nt UMI sequence in the RNA adapter was extracted, and reads were deduplicated after mapping using umi_tools. Mapping was performed using STAR with the Arabidopsis TAIR 10 genome as reference and the following parameters: outFilterMismatchNoverLmax: 0.1; alignIntronMin: 500; alignIntronMax: 1200; outSAMmultNmax: 1. Differentially abundant RNA regions and the region boundaries were identified using DiffSegR^63^. Intron domains were annoated according to the *N. tabacum* domain annotation in the CRW2 database^64^.

### Total plant RNA extraction and northern blotting

Total plant RNA was extracted using GENEzol (Geneaid Biotech) according to the manufacturer instructions with minor modifications. For visual observation of rRNAs, 10 µg of total RNA was loaded onto a 1.5% agarose-formaldehyde gel (20 mM MOPS, pH 7, 8 mM sodium acetate, 1mM EDTA, 1.5% agarose, 6% formaldehyde) and separated by size by running the gel with running buffer (20 mM MOPS, pH 7, 8 mM sodium acetate, 1mM EDTA, 3.7% formaldehyde) at 110 V for 3 hours and stained with ethidium bromide. Quantification of rRNAs was conducted using RNA ScreenTape by a TapeStation device (Agilent).

For RT-qPCR, first-strand cDNA synthesis was performed using the RevertAid First Strand cDNA Synthesis Kit (Thermo Fisher Scientific) according to the manufacturer instructions using a 1:1 mixture of random hexamer primers and oligo(dT)18 primers for cDNA synthesis. After 10-fold dilution, 1 µl cDNA was used in a 10 µl reaction of HOT FIREPol EvaGreen qPCR mix (Solis BioDyne). qPCR reactions were performed using the LightCycler 480 II system (Roche). For quantification of relative abundances of spliced and unspliced transcripts, abundances were normalized to the *AtRCE1* (*AT4G36800*) housekeeping gene^65^ before normalizing to the mean of WT treated with β-estradiol. Primers are listed in **Supplementary Data 1**.

For northern blotting, 10 µg of total RNA was resolved by a 1.5 % agarose-formaldehyde gel and transferred to HyBond-NX Nylon membrane (GE Healthcare) by capillary transfer. The transferred RNA was then cross-linked to the membrane by two rounds of ultraviolet light irradiation (1.2 kJ each). Radiolabelled probes against target transcripts were generated using a Klenow-based labelling kit (Prime-a-Gene, Promega) using synthesized DNA primers as template in the presence of [α-^32^P] dCTP. Probe hybridization, membrane washing, and membrane exposure were conducted as previously described^66^, except that the hybridization and subsequent washing steps were carried out at 60 °C. The probes and the primers used to amplify them are listed in **Supplementary Data 1**.

### Generation of yeast strains

Yeast strains, and constructs and primers used for creating them are detailed in **Supplementary Data 1**. Constructs were made following the modular cloning toolkit for yeast^67^. Plasmids were validated as described for plant plasmids. Yeast transformation, transformant selection via hygromycin resistance or uracil prototrophy, strain purification and confirmation of correct genomic integration by PCR were conducted as previously described^31^.

For expression of YFP-tagged orthologous BEs, the CDS of BEs (without their cTPs, unless cTP prediction was weak; **Supplementary Data 1**) were cloned as part 3a into pYTK001 entry vector^67^ via *Bsm*BI. They were then individually assembled into yeast integrative vectors via *Bsa*I into the pre-assembled backbone vector pBP296 (ref. ^31^), which targeted the intergenic *XII-5* yeast locus^68^, using part plasmids containing the strong constitutive *CPW2* promoter, a C-terminal *eYFP* tag (introducing the same linker before the YFP as in the pUBC-YFP based plant expression vectors), and the *CYC1* terminator. The resulting vectors containing single transcription unit (TU) for BE expression were transformed into the haploid CEN.PK113-11C yeast strain “578”^31^ (*MATa MAL2-8^C^ SUC2 his3Δ malx2 glc3Δ gsy2Δ glg1Δ glg2Δ gph1Δ gsy1::pGAL1-EcglgC-TM-HA-tCYC1 bar1Δ XII-2::pCWP2-mCherry-tUPT7:KanR gdb1::pUBI4-AtSS1-tRPL3*).

For IP experiments in yeast, all final yeast integrative vectors were targeted to the intergenic *XII-2* yeast locus^68^, using either single TU constructs (for control strains expressing only a single test protein) or multi TU constructs (for pair-wise interaction tests; **Supplementary Data 1**. The *AtMatK* CDS (codon optimized for Arabidopsis and containing the edit frequently observed in *AtMatK* RNA^69^, resulting in H236Y conversion) was first cloned as part 3a into pYTK001 via *Bsm*BI, then assembled via *Bsa*I into a single-TU yeast vector resulting in the *P_GAL1_:AtMatK-3xHA:T_CYC1_*TU. For expression of *At*MKIP1-YFP and *At*BE3-YFP without their cTPs, the same TUs as for orthologous BE expression (*P_CWP2_:AtMKIP1-YFP:T_CYC1_*and *P_CWP2_:AtBE3-YFP:T_CYC1_*) were used but targeting them to locus *XII-2*. For expression of *At*MKIP1 fragments, the corresponding regions (AA 120-333 for CBM-MSR; AA 178-333 for MSR; numbers based on the full-length *At*MKIP1 protein including its cTP) were cloned as part 3a into pYTK001, then assembled into equivalent single-TU vectors as the native *At*MKIP1 protein. The ΔMSR-version of *At*MKIP1 was created by amplifying the *At*MKIP1(ΔMSR) fragment from the vector AtMKIP1(ΔMSR)_pUBC-YFP and assembling into *Hind*III pre-digested pBP595 containing *PCWP2:AtMKIP1-YFP:TCYC1* using the NEBuilder HiFi DNA Assembly kit (New England Biolabs). N-terminal (AA 1-381) and C-terminal (AA 357-526) *At*MatK segments were created by deleting the other MatK regions in the pBP662 vector containing the *P_GAL1_:AtMatK-3xHA:T_CYC1_* TU by DNA assembly as above. For pair-wise interaction tests, multi-TU integrative vectors were fused via *Bsm*BI using the preassembled integrative vector pBP492, single-TU vectors and primer-based spacers occupying empty positions. Final single- and multi-TU vectors were *Not*I digested and transformed into the haploid CEN.PK113-11C strain “344.1”^31^ (*MATa MAL2-8^C^ SUC2 his3Δ ura3-52 glc3Δ gsy2Δ glg1Δ glg2Δ*).

### Light microscopy and branching enzyme activity gels in yeast

For light microscopy of iodine stained yeast cells, yeasts were inoculated in complex medium with 2% (w/v) glucose (YPD) until saturation, diluted in complex medium with 2% (w/v) galactose (YPGal, to induce expression of *P_GAL1_*-driven ADPglucose pyrophosphorylase) to an OD of ∼0.3 in Erlenmeyer flasks. After shaking for 5.75 h at 30°C and 260 rpm, cells were harvested by centrifugation, washed with water, flash-frozen in liquid N_2_ and stored at -80°C. Cells were thawn on ice, stained by mixing with Lugol’s solution in a 1:1 ratio, and immediately imaged using an Axio Imager.Z2 light microscope equipped with a 100X oil-immersion objective (Zeiss).

For protein analysis, yeasts were grown in the same way, but the main cultures were harvested already after 3 h shaking in YPGal. Total and soluble protein extracts for immunoblotting were prepared as previously described^31^. Immunoblotting was conducted as described above for plant proteins. Native protein extraction and native PAGE monitoring BE activities (zymograms) were conducted as previously described^22^.

### Immunoprecipitation experiments in yeast

Yeasts were inoculated in YDP, grown for ∼6h at 30°C and 260 rpm, then inoculated to an OD of ∼0.015 in Erlenmeyer flasks and shaken overnight at 30°C and 260 rpm. Cells were harvested by centrifugation and once washed in water. Fresh cell pellets were resuspended in 3.3 volumes of native protein extraction buffer (0.1 M MOPS, pH 7.75, 1% [v/v] Triton X-100, 1 mM EDTA, 4 mM MgCl_2_, protease inhibitor [Complete EDTA-free; Roche]) and 4.3 volumes of glass beads (acid-washed, 425–600 mm diameter) and homogenized by vortexing for 30 min at 4°C. The homogenate was transferred without glass beads to a fresh tube and centrifuged at 16,000 *g* for 5 min at 4°C. Equal amounts of soluble proteins present in the supernatant were mixed with pre-blocked and/or washed magnetic beads. In case of anti-HA beads (Anti-HA Magnetic Beads, Pierce), beads had been pre-blocked by incubation for 30 min at room temperature in TBS-T (Tris-buffered saline, 0.1% [v/v] Tween-20) supplemented with 5% (w/v) bovine serum albumin (BSA), then washed with TBS-T and resuspended in native protein extraction buffer. Magnetic agarose anti-YFP beads (GFP-Trap, ChromoTek) had been prepared by washing them once in wash buffer (50 mM Tris-HCl, pH 8, 0.15 M NaCl, 1 mM DTT, 1% [v/v] Triton X-100) and resuspending them in native protein extraction buffer. The protein-bead mixture was incubated on a rotating wheel at 4°C for 1 h, then washed six times with wash buffer (for anti-YFP beads) or TBS-T (for anti-HA beads), using a magnetic stand for bead recovery. In the last wash step, beads were transferred to a fresh tube to remove proteins adhering to the plastic walls. Proteins bound to the beads were eluted by boiling in SDS-PAGE loading buffer as described for Arabidopsis IP above. Input samples are aliquots of soluble proteins not subjected to immunoprecipitation. Replicates derive from individual main cultures.

### Sequence analyses and protein structure prediction

Starch-branching enzyme protein sequences were identified by BLASTp using *At*MKIP1, *At*BE2, *At*BE3, and rice BEI (a class I BE) as query in multiple genome and transcriptome databases. Sequences were manually curated to remove duplicates and sequences shorter than 200 amino acids very likely constituting fragments. Sequences were aligned by MAFFT (v7) with the E-INS-I algorithm. Phylogenetic analyses were conducted in PhyloBayes MPI (v1.8c) using the “CAT + GTR” substitution model. Two independent runs were performed, each cycling until the sampled tree had stabilized and reached satisfactory convergence (maxdiff <0.3). A convergent tree was calculated from the two runs and the posterior probabilities computed, discarding the first 500 cycles (< 5% of total cycles) of each run as burn-in. The unrooted phylogenetic tree was plotted using Interactive Tree of life (v6).

For protein conservation analysis, protein sequences of each BE class (defined by the phylogenetic tree) were aligned using MAFFT (v7) with the E-INS-I algorithm. Motif logos of regions of interest were constructed using WebLogo 3. BE domains were predicted by SMART, Interpro and a locally installed version of ChloroP. Disordered regions were predicted by AIUPred. The MSR was inferred from the multiple sequence alignment. The conserved regions in the central catalytic domain were defined as previously^30^. RT motifs of *Nt*MatK are annotated as described previously^15^. Protein structures were predicted by AlphaFold2^70^. Molecular graphics and analyses were performed with UCSF ChimeraX. The protein-protein interaction interface from the predicted *At*MatK-*At*MKIP1 dimer was extracted using PDBsum.

## Accession numbers

The Arabidopsis Genome Initiative gene codes associated with this study are: AT3G20440 (*AtMKIP1/AtBE1*), ATCG00040 (*AtMatK*), AT1G07320 (*AtuL4c*), AT5G16715 (*AtValRS2*), AT3G14900 (*AtEMB3120*), AT2G36390 (*AtBE3*). Accession numbers for BE orthologs expressed in yeast are given in **Supplementary Data 1**.

## Statistical analyses

The data shown in **Figs. 6d, 7c-e** and **Supplemental Fig. 10** were first evaluated for normal distribution using the Shapiro-Wilk Test and tested for equal variance using Bartlett’s test. Since the data did not meet the assumptions of normal distribution and equal variance, the non-parametric Kruskal-Wallis test was used for determining statistically significant differences between the groups, followed by a two-sided Dunn’s post hoc multiple comparisons test and Bonferroni *p* value adjustment to identify the specific groups that are significantly different. Statistical analyses of the mass spectrometry and RNA sequencing data is described in the corresponding sections.

## Author contributions

B.P., Y.L., and S.C.Z. designed the research. Y.L. conducted most of the research. Y.G., A.F., M.A., A.G., C.L., M.S., and B.P. collected and analyzed data. Y.L. prepared the figures. B.P. and Y.L. wrote the manuscript. S.C.Z, A.G., R.Z., M.S., and Y.G. edited the manuscript. B.P., S.C.Z., and R.Z. supervised the research and acquired funding. All authors read and approved the manuscript.

## Competing interests

The authors declare no competing interests.

## Acknowledgements

We thank Martha Stadler for excellent technical support, Andrea Ruckle for help with plant cultivation, Christian Schmitz-Linneweber (HU Berlin) for providing us with the *Nt*MatK C+ and C-tobacco lines and Ari Pekka Mähönen (University of Helsinki) for providing us with the pH7m34GW vector. We thank Paolo Nanni and Tobias Kockmann from the Functional Genomics Center Zurich (FGCZ) for the proteomics analyses. AlphaFold2 structure predictions were performed on the ETH Zurich Euler computing cluster. This project has received funding from the European Union’s Horizon 2020 research and innovation programme under the Marie Sklodowska-Curie (MSC) grant agreement No 847585 (to S.C.Z. and B.P.). R.Z. is supported by the Max Planck Society and the DFG grant ZO 302/5-1.

## Notes

### Competing Interest Statement

The authors have declared no competing interest.

## References

1. Germain, A., Hotto, A. M., Barkan, A. & Stern, D. B. RNA processing and decay in plastids. Wiley Interdiscip. Rev. RNA 4, 295–316 (2013).

2. Small, I., Melonek, J., Bohne, A. V., Nickelsen, J. & Schmitz-Linneweber, C. Plant organellar RNA maturation. Plant Cell 35, 1727–1751 (2023).

3. Galej, W. P., Toor, N., Newman, A. J. & Nagai, K. Molecular Mechanism and Evolution of Nuclear Pre-mRNA and Group II Intron Splicing: Insights from Cryo-Electron Microscopy Structures. Chem. Rev. 118, 4156–4176 (2018).

4. Toor, N., Keating, K. S., Taylor, S. D. & Pyle, A. M. Crystal structure of a self-spliced group II intron. Science (80-.). 320, 77–82 (2008).

5. Matsuura, M., Noah, J. W. & Lambowitz, A. M. Mechanism of maturase-promoted group II intron splicing. EMBO J. 20, 7259–7270 (2001).

6. Zhao, C. & Pyle, A. M. The group II intron maturase: a reverse transcriptase and splicing factor go hand in hand. Curr. Opin. Struct. Biol. 47, 30–39 (2017).

7. Cui, X., Matsuura, M., Wang, Q., Ma, H. & Lambowitz, A. M. A group II intron-encoded maturase functions preferentially In Cis and requires both the reverse transcriptase and X domains to promote RNA splicing. J. Mol. Biol. 340, 211–231 (2004).

8. Qu, G. et al. Structure of a group II intron in complex with its reverse transcriptase. Nat. Struct. Mol. Biol. 23, 549–557 (2016).

9. Xu, L., Liu, T., Chung, K. & Pyle, A. M. Structural insights into intron catalysis and dynamics during splicing. Nature 624, 682–688 (2023).

10. Muino, J. M. et al. MatK impacts differential chloroplast translation by limiting spliced tRNA-K(UUU) abundance. Plant J. 1–16 (2024). doi:10.1111/tpj.16945

11. Zoschke, R. et al. An organellar maturase associates with multiple group II introns. Proc. Natl. Acad. Sci. USA 107, 3245–3250 (2010).

12. Lemieux, C., Otis, C. & Turmel, M. Comparative chloroplast genome analyses of streptophyte green algae uncover major structural alterations in the Klebsormidiophyceae, Coleochaetophyceae and Zygnematophyceae. Front. Plant Sci. 7, (2016).

13. Vogel, J. & Börner, T. Lariat formation and a hydrolytic pathway in plant chloroplast group II intron splicing. EMBO J. 21, 3794–3803 (2002).

14. Schmitz-Linneweber, C., Lampe, M. K., Sultan, L. D. & Ostersetzer-Biran, O. Organellar maturases: A window into the evolution of the spliceosome. Biochim. Biophys. Acta 1847, 798–808 (2015).

15. Mohr, G., Perlman, P. S. & Lambowitz, A. M. Evolutionary relationships among group II intron-encoded proteins and identification of a conserved domain that may be related to maturase function. Nucleic Acids Res. 21, 4991–4997 (1993).

16. Hausner, G. et al. Origin and evolution of the chloroplast trnK (matK) intron: A model for evolution of group II intron RNA structures. Mol. Biol. Evol. 23, 380–391 (2006).

17. Till, B., Schmitz-Linneweber, C., Williams-Carrier, R. & Barkan, A. CRS1 is a novel group II intron splicing factor that was derived from a domain of ancient origin. Rna 7, 1227–1238 (2001).

18. Watkins, K. P. et al. A ribonuclease III domain protein functions in group II intron splicing in maize chloroplasts. Plant Cell 19, 2606–2623 (2007).

19. Kroeger, T. S., Watkins, K. P., Friso, G., Van Wijk, K. J. & Barkan, A. A plant-specific RNA-binding domain revealed through analysis of chloroplast group II intron splicing. Proc. Natl. Acad. Sci. U. S. A. 106, 4537–4542 (2009).

20. Pfister, B. & Zeeman, S. C. Formation of starch in plant cells. Cell. Mol. Life Sci. 73, 2781–2807 (2016).

21. Dumez, S. et al. Mutants of Arabidopsis lacking starch branching enzyme II substitute plastidial starch synthesis by cytoplasmic maltose accumulation. Plant Cell 18, 2694– 2709 (2006).

22. Pfister, B. et al. Recreating the synthesis of starch granules in yeast. Elife 5, 1–29 (2016).

23. Wang, X., Xue, L., Sun, J. & Zuo, J. The Arabidopsis *BE1* gene, encoding a putative glycoside hydrolase localized in plastids, plays crucial roles during embryogenesis and carbohydrate metabolism. J. Integr. Plant Biol. 52, 273–288 (2010).

24. Bryant, N., Lloyd, J., Sweeney, C., Myouga, F. & Meinke, D. Identification of nuclear genes encoding chloroplast-localized proteins required for embryo development in Arabidopsis. Plant Physiol. 155, 1678–1689 (2011).

25. Despres, B., Delseny, M. & Devic, M. Partial complementation of embryo defective mutations: A general strategy to elucidate gene function. Plant J. 27, 149–159 (2001).

26. Aryamanesh, N. et al. The Pentatricopeptide Repeat Protein EMB2654 Is Essential for Trans-Splicing of a Chloroplast Small Ribosomal Subunit Transcript. Plant Physiol. 173, 1164–1176 (2017).

27. Yu, H. D. et al. Downregulation of chloroplast RPS1 negatively modulates nuclear heat-responsive expression of HsfA2 and its target genes in Arabidopsis. PLoS Genet. 8, (2012).

28. Han, Y., Sun, F. J., Rosales-Mendoza, S. & Korban, S. S. Three orthologs in rice, Arabidopsis, and *Populus* encoding starch branching enzymes (SBEs) are different from other SBE gene families in plants. Gene 401, 123–130 (2007).

29. Ball, S., Colleoni, C., Cenci, U., Raj, J. N. & Tirtiaux, C. The evolution of glycogen and starch metabolism in eukaryotes gives molecular clues to understand the establishment of plastid endosymbiosis. J. Exp. Bot. 62, 1775–1801 (2011).

30. Tetlow, I. J. & Emes, M. J. A review of starch-branching enzymes and their role in amylopectin biosynthesis. IUBMB Life 66, 546–558 (2014).

31. Pfister, B. et al. Tuning heterologous glucan biosynthesis in yeast to understand and exploit plant starch diversity. BMC Biol. 20, 1–20 (2022).

32. Kuriki, T., Guan, H., Sivak, M. & Preiss, J. Analysis of the active center of branching enzyme II from maize endosperm. J. Protein Chem. 15, 305–313 (1996).

33. Funane, K., Libessart, N., Stewart, D., Michishita, T. & Preiss, J. Analysis of essential histidine residues of maize branching enzymes by chemical modification and site-directed mutagenesis. J. Protein Chem. 17, 579–590 (1998).

34. McBride, A., Ghilagaber, S., Nikolaev, A. & Hardie, D. G. The Glycogen-Binding Domain on the AMPK β Subunit Allows the Kinase to Act as a Glycogen Sensor. Cell Metab. 9, 23–34 (2009).

35. Seung, D., Schreier, T. B., Bürgy, L., Eicke, S. & Zeeman, S. C. Two plastidial coiled-coil proteins are essential for normal starch granule initiation in Arabidopsis. Plant Cell 30, 1523–1542 (2018).

36. Berg, M., Rogers, R., Muralla, R. & Meinke, D. Requirement of aminoacyl-tRNA synthetases for gametogenesis and embryo development in Arabidopsis. Plant J. 44, 866–878 (2005).

37. Asakura, Y., Bayraktar, O. A. & Barkan, A. Two CRM protein subfamilies cooperate in the splicing of group IIB introns in chloroplasts. RNA 14, 2319–2332 (2008).

38. Shinozaki, K. et al. The complete nucleotide sequence of the tobacco chloroplast genome: Its gene organization and expression. EMBO J. 5, 2043–2049 (1986).

39. Kendrick, R., Chotewutmontri, P., Belcher, S. & Barkan, A. Correlated retrograde and developmental regulons implicate multiple retrograde signals as coordinators of chloroplast development in maize. Plant Cell 4897–4919 (2022). doi:10.1093/plcell/koac276

40. Reiter, B. et al. The Arabidopsis protein CGL20 is required for plastid 50S ribosome biogenesis. Plant Physiol. 182, 1222–1238 (2020).

41. Longevialle, F. De et al. The pentatricopeptide repeat gene OTP51 with two LAGLIDADG motifs is required for the cis -splicing of plastid ycf3 intron 2 in Arabidopsis thaliana. Plant J. 56, 157–168 (2008).

42. Wang, C. et al. Rerouting of ribosomal proteins into splicing in plant organelles. Proc. Natl. Acad. Sci. U. S. A. 117, 29979–29987 (2020).

43. Roy, S., Ueda, M., Kadowaki, K. I. & Tsutsumi, N. Different status of the gene for ribosomal protein S16 inthe chloroplast genome during evolution of the genus Arabidopsis and closely related species. Genes Genet. Syst. 85, 319–326 (2010).

44. Zhang, J. et al. Arabidopsis thaliana branching enzyme 1 is essential for amylopectin biosynthesis and cesium tolerance. J. Plant Physiol. 252, 153208 (2020).

45. Anderson, B. M., Krause, K. & Petersen, G. Mitochondrial genomes of two parasitic Cuscuta species lack clear evidence of horizontal gene transfer and retain unusually fragmented ccmFC genes. BMC Genomics 22, 1–17 (2021).

46. Grimaud, F., Rogniaux, H., James, M. G., Myers, A. M. & Planchot, V. Proteome and phosphoproteome analysis of starch granule-associated proteins from normal maize and mutants affected in starch biosynthesis. J. Exp. Bot. 59, 3395–3406 (2008).

47. Helle, S. et al. Proteome Analysis of Potato Starch Reveals the Presence of New Starch Metabolic Proteins as Well as Multiple Protease Inhibitors. 9, 1–14 (2018).

48. Majeran, W. et al. Nucleoid-enriched proteomes in developing plastids and chloroplasts from maize leaves: A new conceptual framework for nucleoid functions. Plant Physiol. 158, 156–189 (2011).

49. Huang, M. et al. Construction of plastid reference proteomes for maize and Arabidopsis and evaluation of their orthologous relationships; The concept of orthoproteomics. J. Proteome Res. 12, 491–504 (2013).

50. Hayashi, M. et al. Bound substrate in the structure of cyanobacterial branching enzyme supports a new mechanistic model. J. Biol. Chem. 292, 5465–5475 (2017).

51. Perron, K., Goldschmidt-Clermont, M. & Rochaix, J. D. A factor related to pseudouridine synthases is required for chloroplast group II intron trans-splicing in Chlamydomonas reinhardtii. EMBO J. 18, 6481–6490 (1999).

52. Jenkins, B. D. & Barkan, A. Recruitment of a peptidyl-tRNA hydrolase as a facilitator of group II intron splicing in chloroplasts. EMBO J. 20, 872–879 (2001).

53. Singh, R. N., Saldanha, R. J., D’Souza, L. M. & Lambowitz, A. M. Binding of a group II intron-encoded reverse transcriptase/maturase to its high affinity intron RNA binding site involves sequence-specific recognition and autoregulates translation. J. Mol. Biol. 318, 287–303 (2002).

54. Hertel, S. et al. Multiple checkpoints for the expression of the chloroplast-encoded splicing factor MatK. Plant Physiol. 163, 1686–1698 (2013).

55. Zoschke, R., Ostersetzer, O., Börner, T. & Schmitz-Linneweber, C. Analysis of the regulation of MatK gene expression. Endocytobiosis Cell Res. 19, 127–135 (2009).

56. Paukstelis, P. J., Chen, J., Chase, E., Lambowitz, A. M. & Golden, B. L. Structure of a tyrosyl-tRNA synthetase splicing factor bound to a group I intron RNA. Nature 451, 94– 98 (2008).

57. Rho, S. B., Lincecum, T. L. & Martinis, S. A. An inserted region of leucyl-tRNA synthetase plays a critical role in group I intron splicing. EMBO J. 21, 6874–6881 (2002).

58. Grefen, C. et al. A ubiquitin-10 promoter-based vector set for fluorescent protein tagging facilitates temporal stability and native protein distribution in transient and stable expression studies. Plant J. 64, 355–365 (2010).

59. Siligato, R. et al. Multisite gateway-compatible cell type-specific gene-inducible system for plants. Plant Physiol. 170, 627–641 (2016).

60. Abt, M. R. et al. STARCH SYNTHASE5, a noncanonical starch synthase-like protein, promotes starch granule initiation in Arabidopsis. Plant Cell 32, 2543–2565 (2020).

61. Barkan, A. Genome-Wide Analysis of RNA–Protein Interactions in Plants. in Plant Systems Biology. Methods in Molecular Biology (ed. Belostotsky, D.) 553, 13–37 (Humana Press, 2009).

62. Ting, M. K. Y. et al. Optimization of ribosome profiling in plants including structural analysis of rRNA fragments. Plant Methods 20, (2024).

63. Liehrmann, A. et al. DiffSegR: an RNA-seq data driven method for differential expression analysis using changepoint detection. NAR Genomics Bioinforma. 5, 1–12 (2023).

64. Cannone, J. J. et al. The Comparative RNA Web (CRW) Site: An online database of comparative sequence and structure information for ribosomal, intron, and other RNAs. BMC Bioinformatics 3, 1–31 (2002).

65. Kindgren, P. et al. The plastid redox insensitive 2 mutant of Arabidopsis is impaired in PEP activity and high light-dependent plastid redox signalling to the nucleus. Plant J. 70, 279–291 (2012).

66. Devers, E. A. et al. Movement and differential consumption of short interfering RNA duplexes underlie mobile RNA interference. Nat. Plants 6, 789–799 (2020).

67. Lee, M. E., DeLoache, W. C., Cervantes, B. & Dueber, J. E. A highly characterized yeast toolkit for modular, multipart assembly. ACS Synth. Biol. 4, 975–986 (2015).

68. Mikkelsen, M. D. et al. Microbial production of indolylglucosinolate through engineering of a multi-gene pathway in a versatile yeast expression platform. Metab. Eng. 14, 104– 111 (2012).

69. Rodrigues, N. F., Christoff, A. P., da Fonseca, G. C., Kulcheski, F. R. & Margis, R. Unveiling Chloroplast RNA Editing Events Using Next Generation Small RNA Sequencing Data. Front. Plant Sci. 8, (2017).

70. Jumper, J. et al. Highly accurate protein structure prediction with AlphaFold. Nature 596, 583–589 (2021).

