## Supplementary Figures and Tables for "Maturase K forms a plastidial splicing complex with a neofunctionalized branching enzyme"

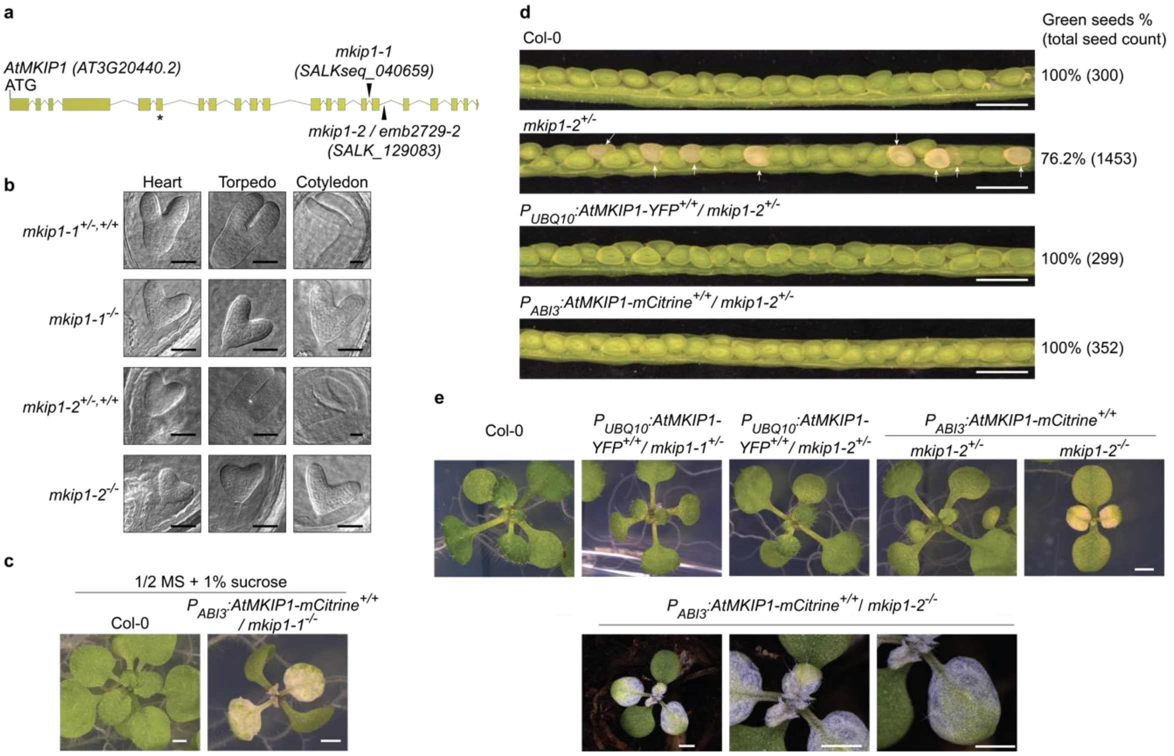


**Supplementary Fig. 1. The embryo defective phenotype of Arabidopsis *mkip1* mutants and their complementation by full-length *AtMKIP1* constructs.**

**a** Genomic structure of the *AtMKIP1* gene. Exons are represented by green boxes and introns by lines. T-DNA insertion sites of the *mkip1* alleles used in this study are indicated by black triangles. Exon 6 (indicated by an asterisk) is absent in splice form 1 but is present in the primary splice form 2 (AT3G20440.2) used for complementation constructs.

**b** Micrographs of wild-type and mutant embryos from siliques of heterozygous *mkip1-1* or *mkip1-2* mother plants at different stages of embryo development. The indicated putative genotypes were inferred from the phenotypes. Scale bars: 50 µm.

**c** Photographs of 20-day-old seedlings grown on ½-strength MS plates containing 1% sucrose. Scale bars: 1 mm.

**d** Opened siliques from mother plants with the indicated genotypes. Seeds with white embryos are indicated by arrows. Percentages indicate the fraction of green seeds of total seeds (number of seeds analyzed in parenthesis). Scale bars: 1 mm.

**e** Photographs of *mkip1* mutant seedlings rescued by expression of *At*MKIP1 under the constitutive *UBIQUITIN10* ($P_{UBQ10}$) or seed-specific *ABI3* ($P_{ABI3}$) promoter. Top panel, photographs of 16-day-old seedlings grown on ½-strength MS plates. Bottom panel, light micrographs of a representative 24-day-old soil-grown *mkip1-2*$^{-/-}$ seedling rescued by the $P_{ABI3}$:*AtMKIP1-mCitrine* construct. Scale bars: 1 mm.

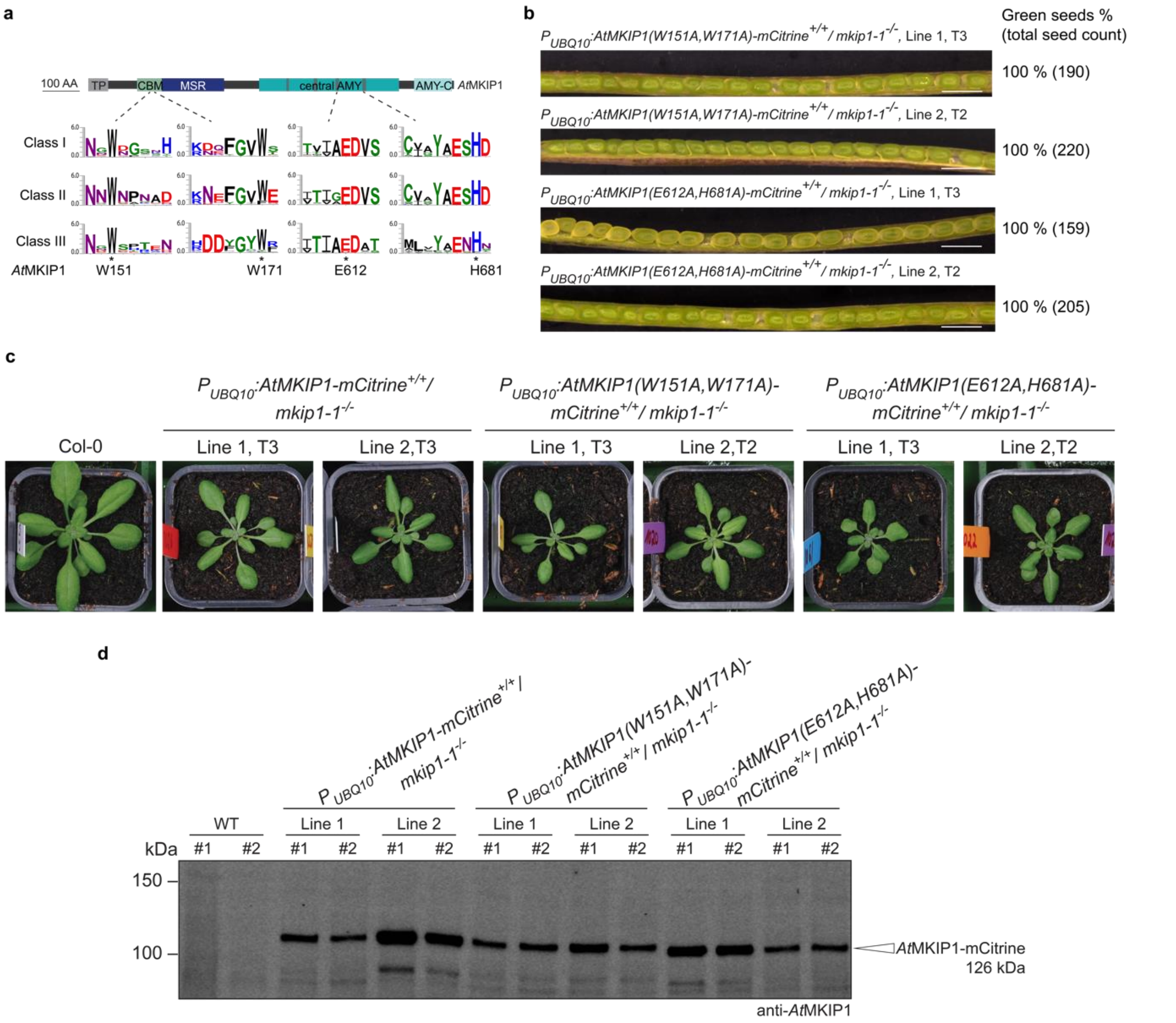


**Supplementary Fig. 2. Complementation of the *mkip1-1* mutant phenotype by *At*MKIP1 variants with point mutations in residues conserved among branching enzymes (BEs).**
**a** WebLogos showing conserved amino acid residues in the carbohydrate-binding module (CBM) and central catalytic domain (central AMY) based on the sequence alignment of BEs from seed plant species (gymnosperms and angiosperms) used for the phylogenetic tree in **Fig. 2a**. Amino acids in *At*MKIP1 that were mutated to alanines are indicated.
**b** Opened siliques from *mkip1-1*$^{+/-}$ mother plants expressing mCitrine-tagged *At*MKIP1 variants. Percentages indicate the proportion of green seeds among total seeds (number of total seeds assessed in parentheses). Two individual lines per construct were evaluated. Scale bars: 1 mm.
**c** Photographs of four-week-old wild-type (Col-0) and *mkip1-1*$^{-/-}$ plants complemented with the indicated wild-type or mutant *At*MKIP1-mCitrine constructs. Representative plants of two individual lines per construct are shown.
**d** Immunoblot detection of the *At*MKIP1 versions introduced by the complementation constructs. Two plants per line were analyzed. Total protein was extracted from a four-week-old rosette leaf and loaded on an equal leaf area basis. Native *At*MKIP1 (98 kDa) in Col-0 could not be detected due to insufficient antibody sensitivity. Source data are provided in the Source Data file.

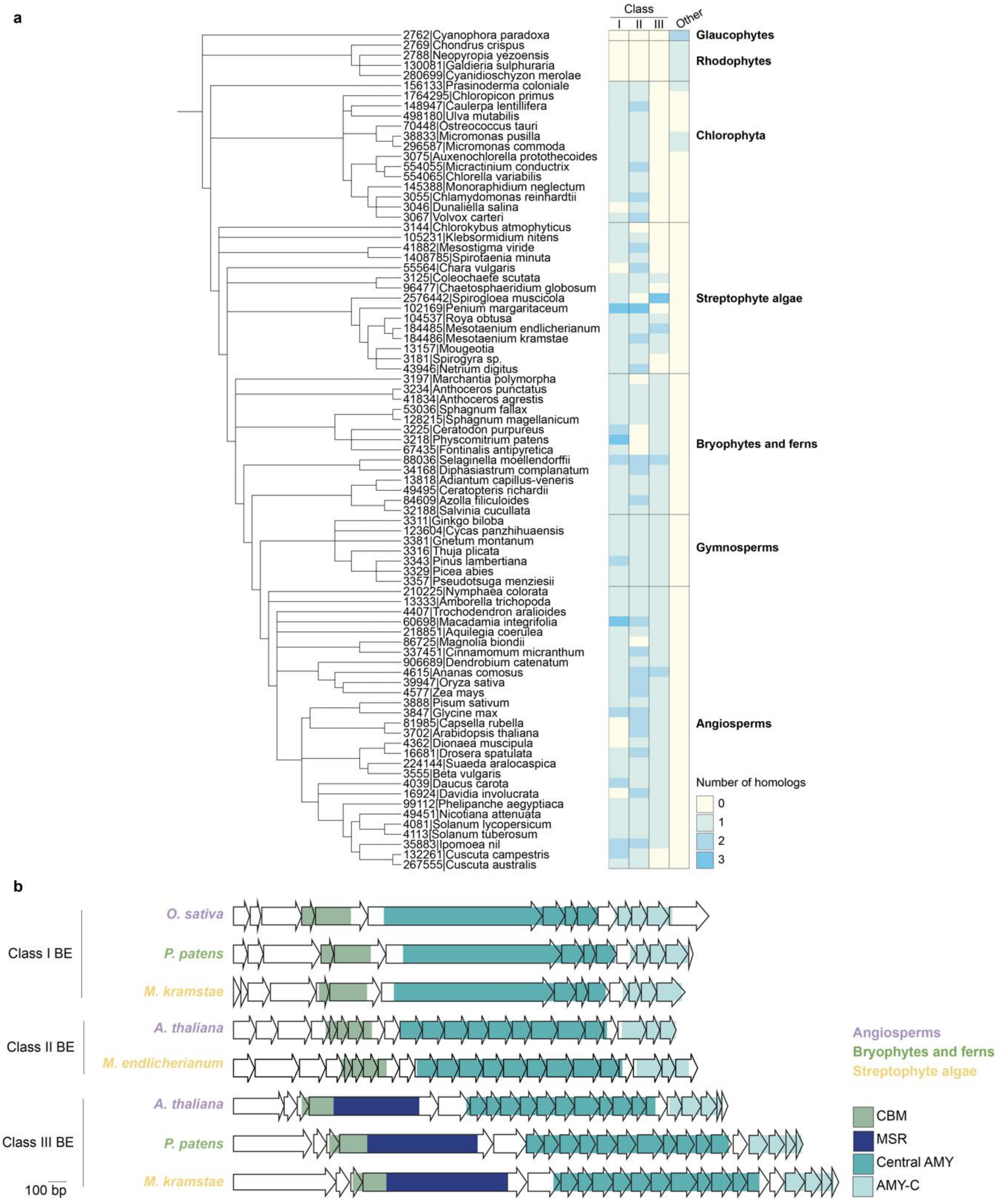


**Supplementary Fig. 3. *BE* gene number and structure in *Archaeplastida*.**

**a** Phylogenetic tree (left) of the species presented in **Fig. 2a** showing the number of genes from each BE class (right). BE classes are defined according to the phylogeny presented in **Fig. 2a**, with „other“ representing the algal BEs that do not fall into classes I, II or III. The species tree follows NCBI taxonomy.

**b** Schematic representation of the exon structure of genes belonging to different BE classes from an angiosperm (*Arabidopsis thaliana* or *Oryza sativa*), a moss (*Physcomitrium patens*) and a streptophyte alga (*Mesotaenium kramstae* or *Mesotaenium endlicherianum*). Each arrow indicates an individual exon. Colors indicate the corresponding encoded protein domains.

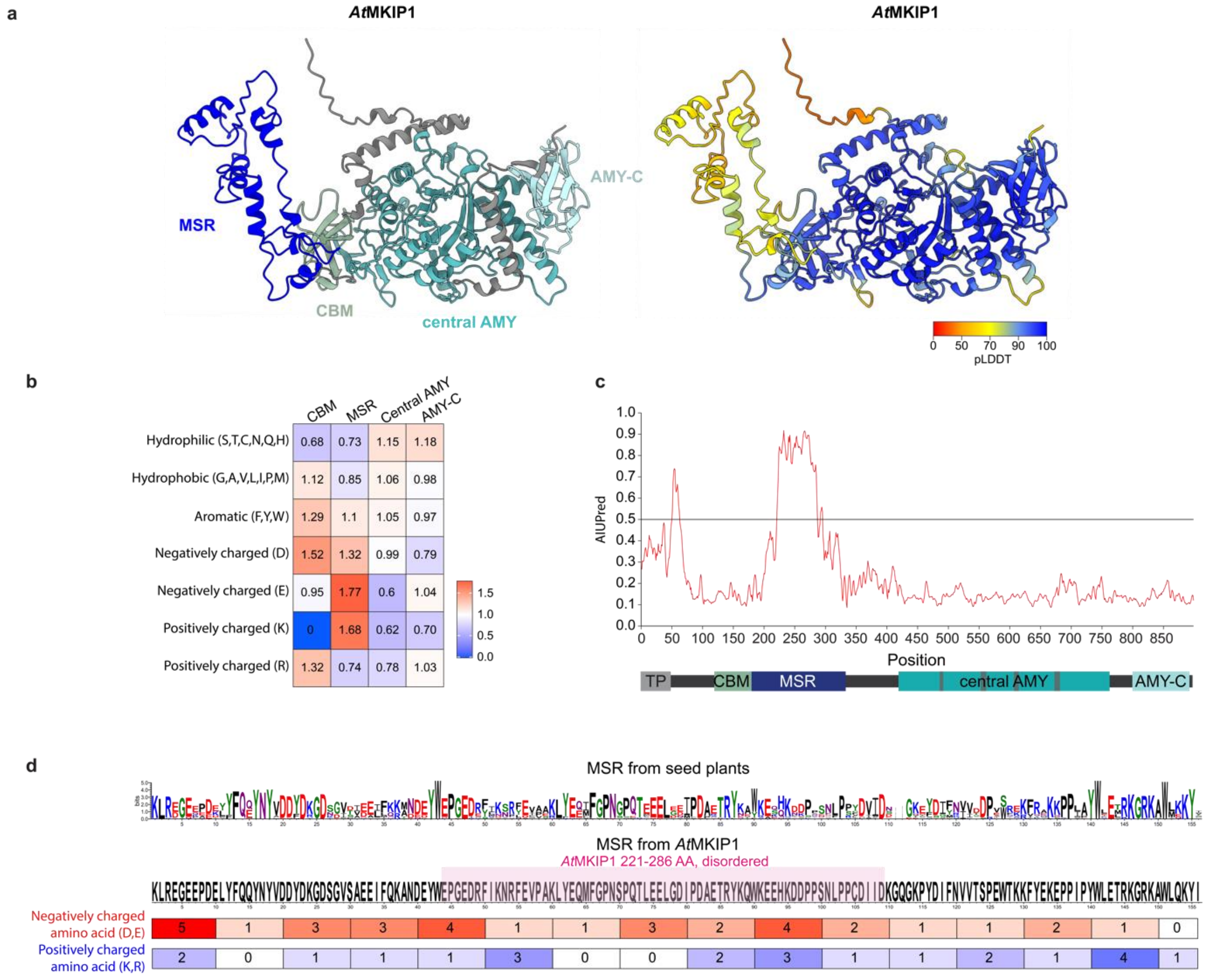


**Supplementary Fig. 4. Amino acid composition and structure prediction of the MKIP1-specific region (MSR).**

**a** AlphaFold2 prediction of *At*MKIP1 without its chloroplast transit peptide. Structure colored by domain (left panel) or local structural confidence (predicted local distance difference test, pLDDT, where red and blue indicate low and high confidence, respectively; right panel). *At*MKIP1 regions are defined as in **Fig. 2b**.

**b** Amino acid composition of *At*MKIP1 regions. The percentage of each amino acid category within each region was normalized to the percentage of that amino acid category in the full-length *At*MKIP1 protein.

**c** Prediction of disordered regions in *At*MKIP1 by AIUPred. A region with a score >0.5 is generally considered to be disordered.

**d** Top panel: WebLogo showing the amino acid conservation in the MSR domain based on the sequence alignment of class III BEs from seed plant species (gymnosperms and angiosperms) used for the phylogenetic tree in **Fig. 2a**. The disordered region predicted by AIUPred is highlighted in pink. Bottom panel: Heatmap of the number of positively and negatively charged amino acids in each ten-amino-acid window within the MSR of *At*MKIP1.

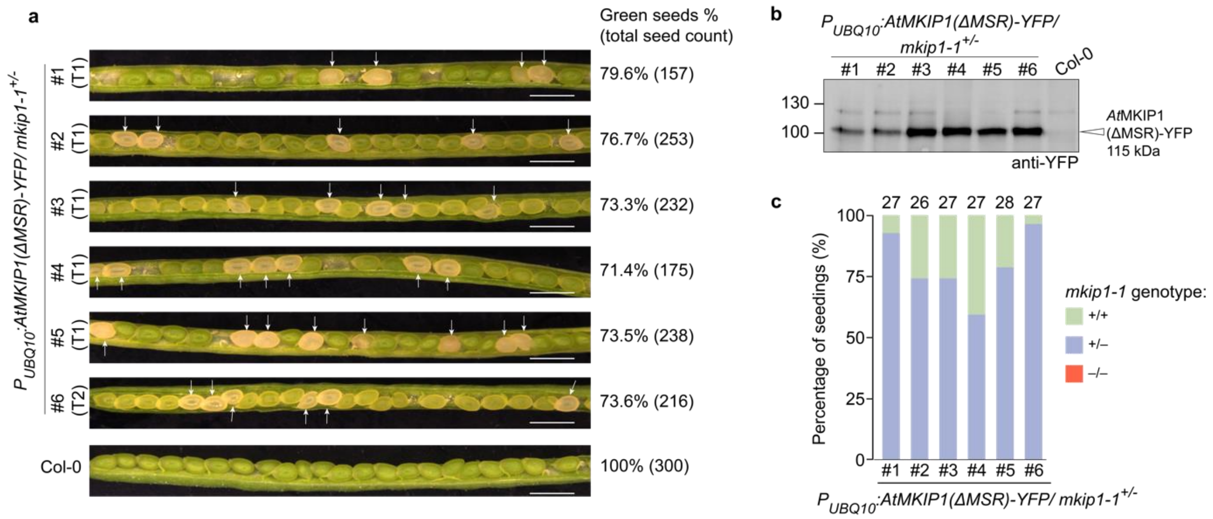


**Supplementary Fig. 5. *At*MKIP1 lacking its MKIP1-specific region (MSR) cannot complement the embryo defects of *mkip1-1*.**

**a** Opened siliques from the wild type (Col-0) and *mkip1-1*$^{+/-}$ transformed with $P_{UBQ10}$:*AtMKIP1(ΔMSR)-YFP, in which the MSR (*amino acids 178-333) of *At*MKIP1 is deleted. White seeds are indicated by arrows. Percentages indicate the proportion of green seeds in total seeds (number of total seeds assessed in parentheses). Scale bars: 1 mm.

**b** Immunoblot analysis of YFP-tagged protein from soluble leaf proteins extracted from 6-week-old rosettes of the mother plants shown in (**a**). Samples were loaded on an equal leaf area basis.

**c** Genotyping of the progeny seedlings from the heterozygous *mkip1-1*$^{+/-}$ shown in (**a**) for the *mkip1-1* T-DNA insertion. The numbers above the bar indicate the total number of seedlings genotyped in each line. No homozygous *mkip1-1*$^{-/-}$ mutant could be identified for any line.

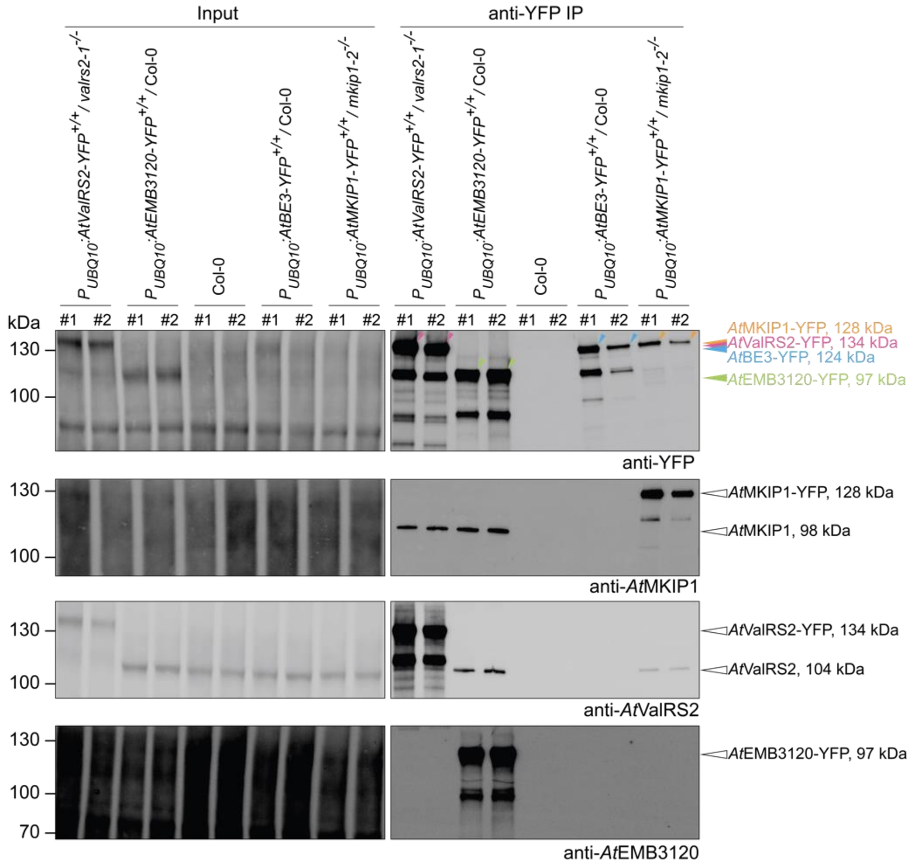


**Supplementary Fig. 6. Reciprocal anti-YFP immunoprecipitation (IP) experiments in Arabidopsis using the YFP-tagged interaction partners of *At*MKIP1 as baits.**
Soluble protein extracted from 10-day-old Arabidopsis seedlings of the indicated genotypes was subjected to anti-YFP IP, using seedlings expressing *At*MKIP1-YFP or *At*BE3-YFP, or no YFP-tagged protein (wild type, Col-0) as controls. Two replicates consisting of ~250 mg (fresh weight) of pooled seedlings each were analyzed for each line. Soluble extracts were analyzed before IP (Input) and after anti-YFP IP. The predicted molecular weights of the proteins refer to the mature proteoforms without their chloroplast transit peptides. The higher apparent molecular weight of *At*EMB3120-YFP is likely due to its high content of negatively charged residues, which often reduce migration velocity in SDS-PAGE. While *At*EMB3120-YFP co-precipitated endogenous *At*MKIP1, *At*MKIP1-YFP did not detectably co-precipitate endogenous *At*EMB3120 in this set of experiments, presumably due to the lower amounts of input material compared to the experiments shown in **Fig. 3b** and the limited sensitivity of the anti-*At*EMB3120 antibody.

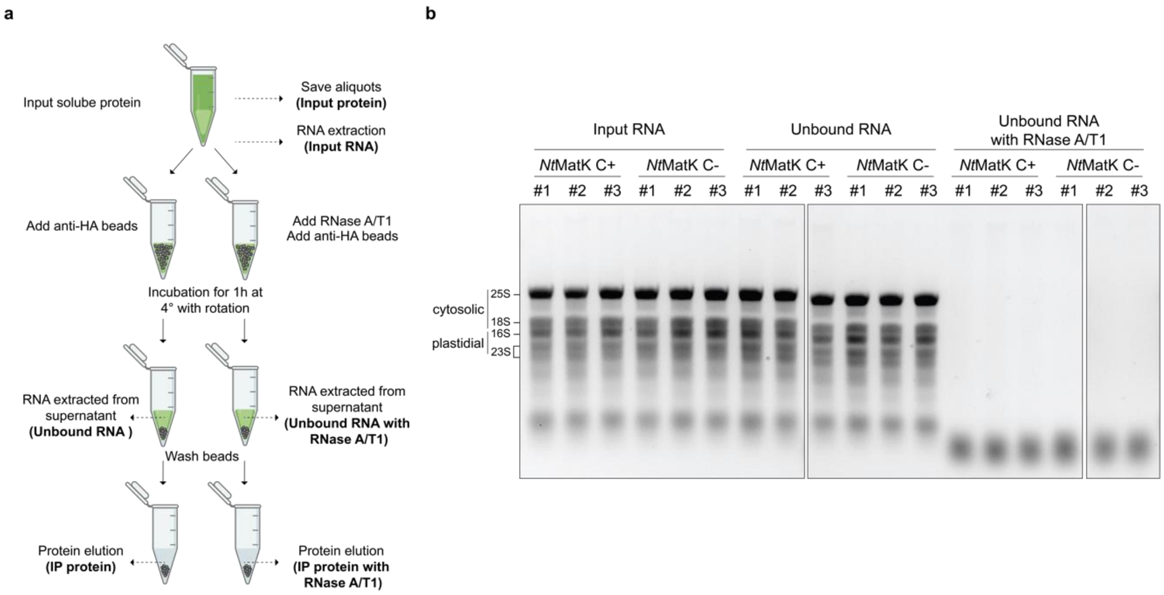


**Supplementary Fig. 7. RNA integrity analysis after RNase A/T1 treatment during the tobacco immunoprecipitation (IP) experiment shown in Fig. 3d.**

**a** Schematic diagram of the IP procedure with or without RNase A/T1 treatment during the bead incubation step. RNA samples collected for analysis are shown in bold.

**b** Gel electrophoresis of RNA isolated during the MatK-HA IP experiment outlined in (**a**). Two µg of RNA were loaded onto a 1.5% agarose gel and visualized by ethidium bromide staining. *Nt*MatK C+ expresses *Nt*MatK with a C-terminal 3xHA tag. *Nt*MatK C- is a control line containing the *aadA* marker gene but no *matK* modification.

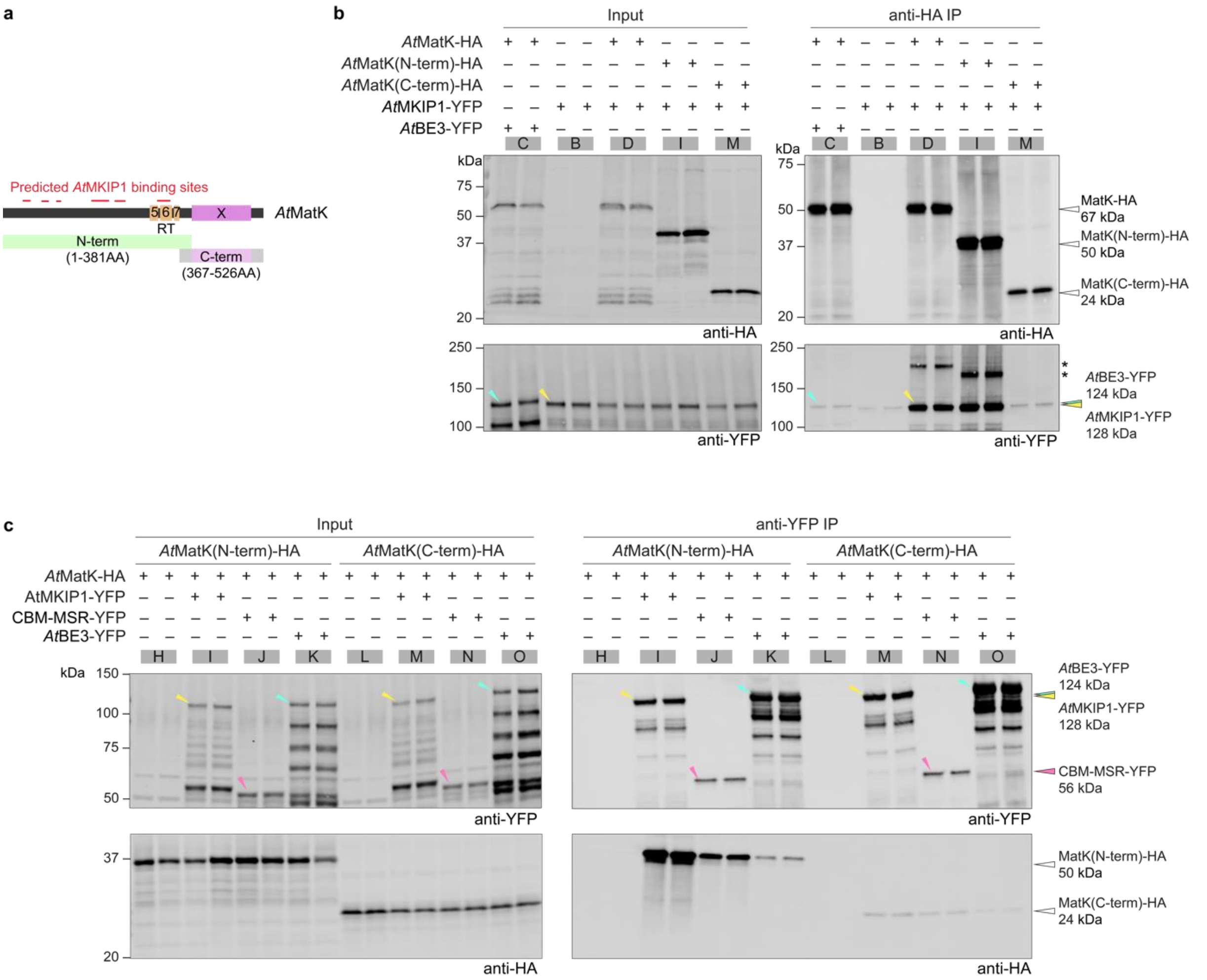


**Supplementary Fig. 8. Reciprocal immunoprecipitation (IP) experiments showing the interaction of *At*MatK-HA and *At*MKIP1-YFP in yeast.**

**a** Schematics of *At*MatK and the fragments used for testing its interaction with *At*MKIP1. Colored boxes represent the reverse transcriptase motifs (RT5-7, *yellow*) and the X domain (*purple*).

**b** Soluble protein extracts from yeast strains expressing the corresponding protein (+) or not (-) were analyzed before IP (input) and after anti-HA IP by immunoblotting using the indicated antibodies. Two replicate cultures were assessed for each strain. The bands marked with asterisks are of unknown nature but fit the predicted molecular weights of *At*MKIP1-*At*MatK dimers, indicating that a fraction of co-precipitated dimers may be resistant to boiling in SDS.

**c** IP experiments performed as in (**b**) but using yeast strains expressing the N- and C-terminal *At*MatK-HA segments shown in (**a**). This figure shows the full IP experiment presented in **Fig. 4e**.

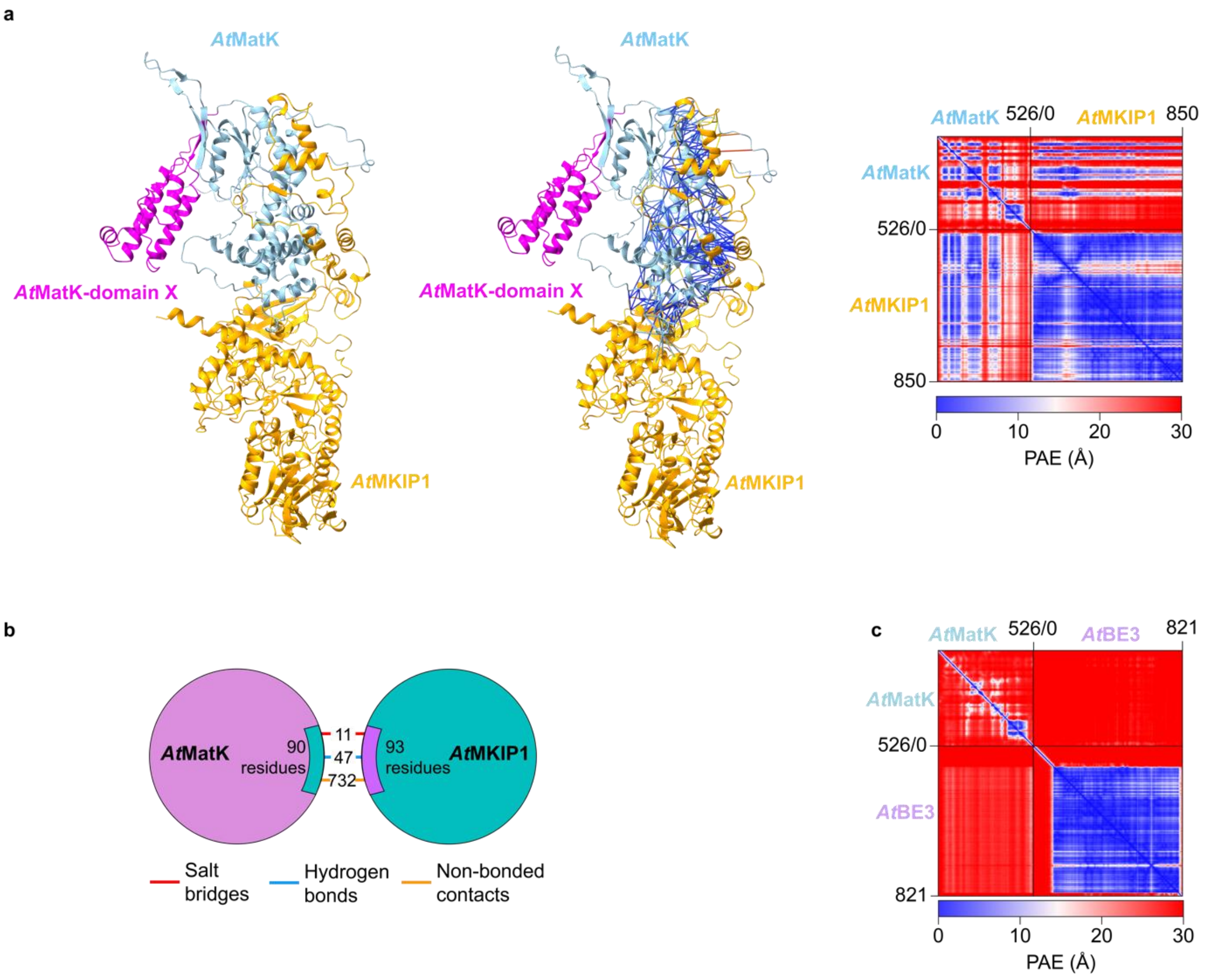


**Supplementary Fig. 9. *At*MatK and *At*MKIP1 are predicted to interact through a large interaction platform that forms multiple intermolecular contacts.**

**a** AlphaFold2 prediction of the protein dimer formed between *At*MatK (in blue, with domain X in magenta) and *At*MKIP1 (in orange). Interprotein residue contacts within 4 Å are marked with lines and colored by their predicted aligned error (PAE), i.e. blue lines indicate high confidence contact sites between *At*MatK and *At*MKIP1.

**b** The protein-protein interaction interface extracted from the predicted *At*MatK-*At*MKIP1 dimer using PDBsum.

**c** Low confidence AlphaFold2 prediction of the protein dimer between *At*MatK and *At*BE3, as indicated by the red coloring in the intermolecular regions in the predicted aligned error (PAE) plot.

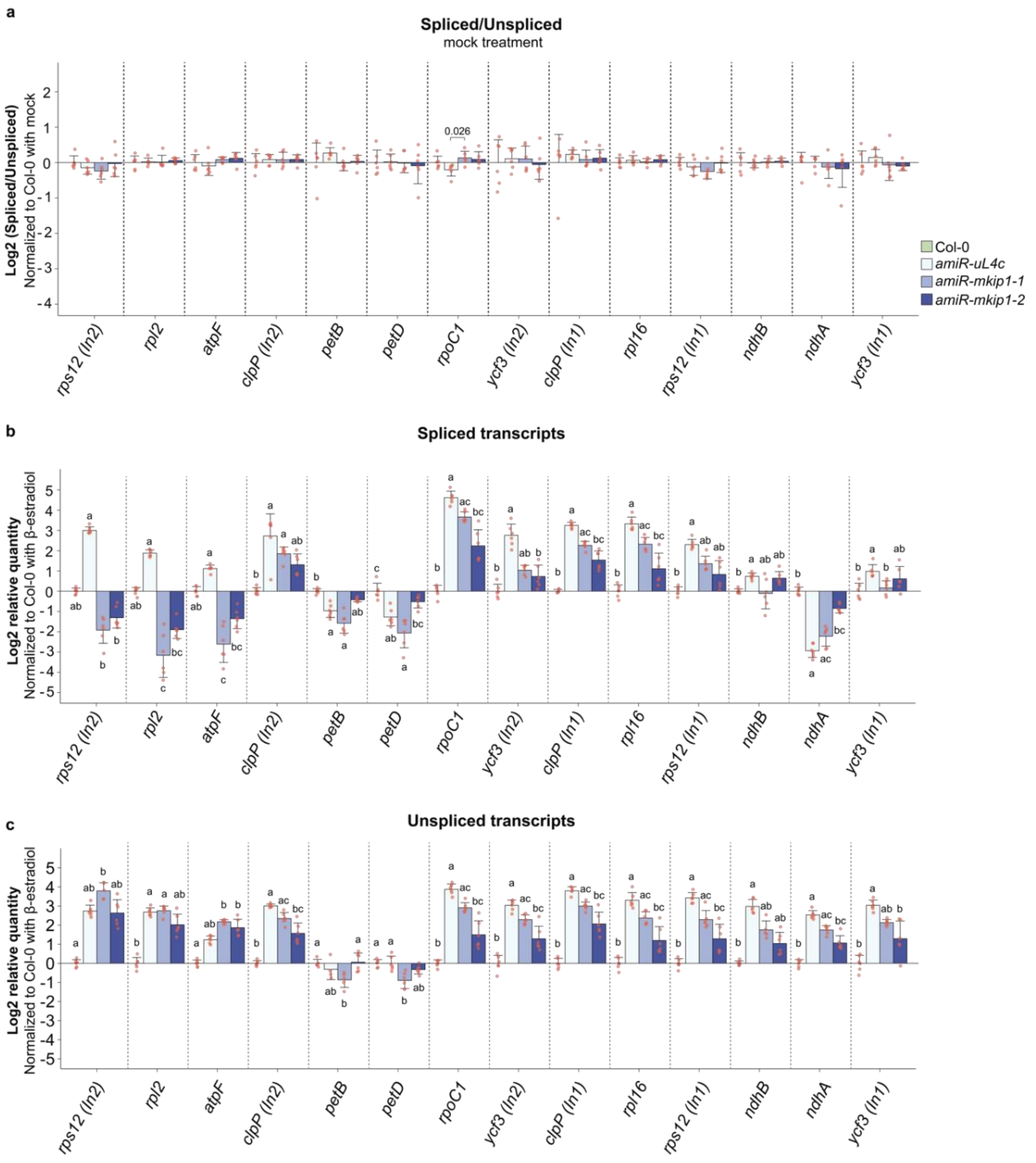


**Supplementary Fig. 10. Quantification of spliced and unspliced mRNAs in *AtMKIP1* silenced and mock-treated plants by RT-qPCR.**

**a** Intron splicing efficiencies in mock-treated plants. Efficiencies were assessed as **Fig. 7d** but normalized to the mean of mock-treated wild type (Col-0). Shown are means ± s.d. ($n$=6 plants). Red dots show individual data points. Statistical significance was evaluated using the Kruskal-Wallis test followed by a two-sided Dunn's post hoc multiple comparisons test and Bonferroni $p$ value adjustment. For statistically significant differences between means ($p$<0.05), the $p$-values are provided above the brackets.

**b-c** Relative abundances of spliced (**b**) and unspliced (**c**) transcripts in β-estradiol-treated plants. Abundances were first normalized to those of the housekeeping gene *RCE1*, then normalized to the mean of β-estradiol-treated wild type (Col-0). Shown are means ± s.d. ($n$=6 plants). Red dots show individual data points. The underlying data are the same as in **Fig. 7d-e**. Statistical significance was evaluated as in (**a**). Different letters indicate statistically significant differences ($p$<0.05) between the means.

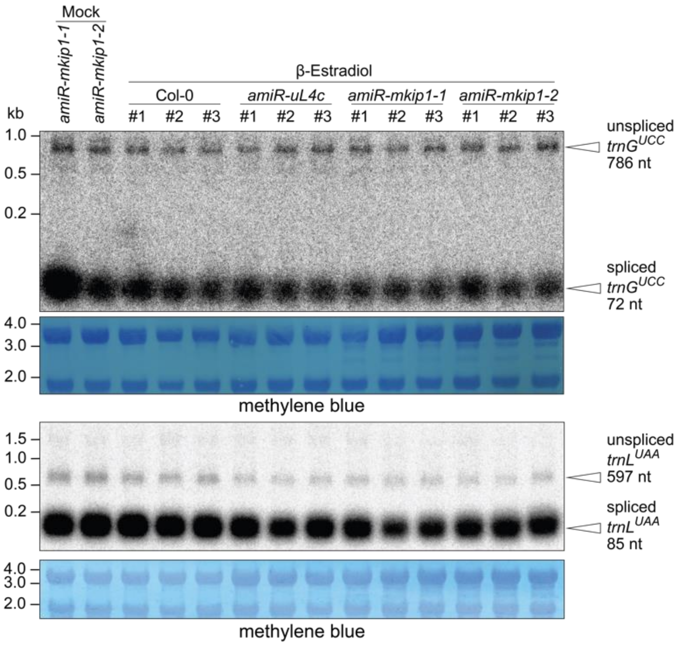


**Supplementary Fig. 11. Northern blot analysis of the splicing of *trnG*$^{UCC}$ and *trnL*$^{UAA}$ introns upon induced *AtMKIP1* silencing.**

β-Estradiol treatment and sampling of newly emerged leaves were conducted as described for **Fig. 6**. Total RNA was separated on denaturing agarose gels, transferred to nylon membranes, and hybridized with probes for *trnG*$^{UCC}$ exon 2 (top panel) or *trnL*$^{UAA}$ exon 1 (third panel). Methylene blue is a total RNA stain used as a loading control. Three estradiol-treated plants were analyzed per line. Spliced and unspliced transcripts (with their calculated sizes) are indicated by arrows. Data were used to calculate the splicing efficiencies shown in **Fig. 7**. Col-0, wild type.

| | | | | Quantification (MS1) | | Razor spectral counts (MS2) | | | | | |
|---|---|---|---|---|---|---|---|---|---|---|---|
| | | | | | | *Nt*MatK C+ | | | *Nt*MatK C- | | |
| **UniProt ID** | **Description** | **cTP** | **Closest Arabidopsis ortholog** | **log2 FC** | **Adj. *p*-value** | **#1** | **#2** | **#3** | **#1** | **#2** | **#3** |
| A0A1S4C7U0_TOBAC | *Nt*EMB3120a | yes | *At*EMB3120 (AT3G14900) | 8.32 | 0.0057 | 157 | 164 | 162 | 0 | 10 | 0 |
| A0A140G1P3_TOBAC | *Nt*MatK | n.a. | *At*MatK (ATCG00040) | 7.83 | 0.0007 | 115 | 155 | 137 | 0 | 8 | 0 |
| A0A1S4BH73_TOBAC | *Nt*ValRS2b | yes | *At*ValRS2 (AT5G16715) | 7.77 | 0.0007 | 196 | 223 | 228 | 0 | 10 | 1 |
| A0A1U7Y1B8_NICSY | *Nt*MKIP1a | yes | *At*MKIP1 (AT3G20440) | 7.23 | 0.0003 | 203 | 204 | 239 | 1 | 16 | 1 |
| A0A1S3ZFZ9_TOBAC | *Nt*EMB3120b | yes | *At*EMB3120 (AT3G14900) | 7.12 | 0.0001 | 9 | 9 | 11 | 0 | 0 | 0 |
| A0A1S3Z171_TOBAC | *Nt*MKIP1b-1 | yes | *At*MKIP1 (AT3G20440) | 6.99 | 0.0002 | 5 | 8 | 6 | 0 | 0 | 0 |
| A0A1S4A7L4_TOBAC | *Nt*MKIP1b-2 | no | *At*MKIP1 (AT3G20440) | 5.91 | 0.0024 | 49 | 60 | 52 | 0 | 1 | 0 |
| A0A1S4AQM5_TOBAC | CRM-domain containing factor | yes | AT3G18390 | 2.61 | 0.0003 | 7 | 8 | 6 | 0 | 0 | 0 |
| A0A1S4DJ40_TOBAC | Ribonuclease III domain protein RNC1 | yes | *At*RNC1 (AT4G37510) | 2.25 | 0.0003 | 6 | 6 | 6 | 1 | 1 | 1 |
| A0A1S3XJ11_TOBAC | Co-chaperone GrpE family protein | yes | *At*CGE2 (AT1G36390) | 1.99 | 0.0028 | 5 | 5 | 4 | 0 | 0 | 1 |
| A0A1S3XZX4_TOBAC | tRNA ribonuclease (tRNase) Z | yes | *At*TRZ2 (AT2G04530) | 1.90 | 0.2241 | 8 | 9 | 5 | 0 | 1 | 0 |
| A0A1S3XQS5_TOBAC | Nuclear pore complex protein NUP98A-like | n.a. | *At*DRA2 (AT1G10390) | 1.80 | 0.0075 | 0 | 4 | 2 | 2 | 1 | 1 |
| A0A1S4DSL7_TOBAC | Chaperonin-60 alpha | yes | *At*CPNA1 (AT2G28000) | 1.77 | 0.0003 | 26 | 26 | 29 | 16 | 17 | 16 |
| A0A1S4AHG8_TOBAC | Chaperonin-60 beta | yes | *At*CPNB2 (AT3G13470) | 1.72 | 0.0002 | 37 | 44 | 47 | 22 | 21 | 29 |
| A0A1S4BNP6_TOBAC | Chaperonin-60 beta | yes | *At*CPNB2 (AT3G13470) | 1.71 | 0.0007 | 2 | 2 | 3 | 0 | 0 | 1 |
| A0A1S3ZEH5_TOBAC | Ankyrin-repeat protein | yes | *At*STT2 (AT5G66055) | 1.68 | 0.0034 | 0 | 2 | 2 | 0 | 0 | 0 |
| A0A1S4A1Q4_TOBAC | Chaperonin-60 beta | yes | *At*CPNB1 (AT1G55490) | 1.52 | 0.0003 | 8 | 8 | 9 | 2 | 2 | 2 |
| A0A1S4CB32_TOBAC | 70 kDa heat shock protein | yes | *At*cpHsc70-2 (AT5G49910) | 1.50 | 0.0002 | 17 | 23 | 21 | 11 | 12 | 12 |
| A0A1S3ZAW0_TOBAC | Heat shock protein 90 | yes | *At*HSP90C (AT2G04030) | 1.19 | 0.0022 | 5 | 6 | 6 | 1 | 0 | 1 |

**Supplementary Table 1. List of proteins that are at least two-fold enriched in the *Nt*MatK C+ anti-HA immunoprecipitation presented in Fig. 3e.**

Protein quantification for calculation of fold changes (FC) and adjusted (adj.) *p* values base on precursor (MS1) intensities. MS2 razor spectral counts are shown in addition to illustrate peptide abundances. Note, however, that in the MS2 counts each spectrum is ultimately assigned to only one razor (or leading) protein, even if it would match to multiple proteins. For highly similar proteins (such as the "a" and "b" isoforms of *Nt*MKIP1, *Nt*EMB3120 or *Nt*ValRS2), where most peptides can be mapped to both homoeologs, the true distribution of spectral counts assigned to either homoeolog is likely to be more balanced.

The presence of chloroplast transit peptides (cTPs) in the tobacco proteins was predicted by ChloroP. The closest Arabidopsis homolog represents the top BLASTp hit against the Arabidopsis proteome, with the associated locus indicated in parentheses. Arabidopsis orthologs predicted or shown to be plastid-localized (SUBA5 consensus localization) are shown in green.

Since *N. tabacum* is an allotetraploid species resulting from a hybridization between *N. sylvestris* and *N. tomentosiformis*, the isoforms of *Nt*MKIP1, *Nt*ValRS2 and *Nt*EMB3120 were designated as "a" and "b" according to their probable origin (based on amino acid identity) from *N. sylvestris* and *N. tomentosiformis*, respectively. The origin of the single plastidial *NtMatK* gene is unclear. *Nt*MKIP1a is the full-length MKIP1 from *N. sylvestris*. *Nt*MKIP1a is not annotated in the *N. tabacum* UniProt reference proteome (UP000084051) but shares 100% amino acid identity with *Nt*MKIP1a that we cloned from *N. tabacum* cDNA. We thus consider *Nt*MKIP1a to be a genuine, full-length *Nt*MKIP1.

*Nt*MKIP1b-1 and *Nt*MKIP1b-2 (annotated in the UniProt *N. tabacum* reference proteome) represent N- and C-terminal halves of *Nt*MKIP1 with short extensions. Whether the fragmentation of *Nt*MKIP1b is a sequencing/annotation artifact is currently unclear.

The remaining proteins are likely to be correctly annotated, full-length proteins except for the NUP98A-like nuclear pore complex protein (A0A1S3XQS5_TOBAC), which likely stems from an incomplete sequence resulting in an N-terminal truncation and was hence excluded from the cTP prediction. Only one *Nt*ValRS2 isoform (*Nt*ValRS2b) was quantified because *Nt*ValRS2a and *Nt*ValRS2b are highly similar and the number of peptides uniquely assigned to *Nt*ValRS2a was insufficient for quantification. n.a., not applicable.
